# A co-transcriptional mechanism for tightly controlling RNA homeostasis in yeast

**DOI:** 10.1101/2024.10.29.620883

**Authors:** Sofia Esteban-Serna, Tove Widén, Mags Gwynne, Iseabail Farquhar, Michael R Duchen, Peter S Swain, Sander Granneman

## Abstract

Transcription termination by the Nrd1-Nab3-Sen1 (NNS) complex is key in repressing pervasive transcription in *Saccharomyces cerevisiae*. Counterintuitively, during starvation, multiple mRNAs that are upregulated are also increasingly bound and prematurely terminated and degraded via NNS. Here we demonstrate that this NNS-mediated attenuation is important for controlling the expression and protein concentration of an evolutionarily conserved mitochondrial transporter, Pic2. Strikingly, we find that even a modest increase in Pic2 protein levels caused by defective NNS regulation has major phenotypical consequences, increasing cell volume and intracellular stress, prolonging cell cycle and decreasing growth rate. Disrupting Nab3 binding to *PIC2* globally redistributed Nrd1 binding, changing the levels of other NNS-regulated transcripts. We propose that imbalances in the availability of the subunits constituting the NNS complex underlie the cell volume and cycle anomalies. Collectively our results illustrate that even subtle changes in how RNA-binding proteins interact with a single RNA substrate can cause global defects and they emphasise the crucial role of the NNS complex in preserving microbial fitness during stress.

**Highlights:**

- NNS regulates the expression and protein concentration of a stress-response protein-coding gene (*PIC2*), improving cell fitness and adaptability to environmental challenges.
- Creating an imbalance in RNA binding of Nab3 and Nrd1 for *PIC2* mRNA disturbs the homeostasis of co-regulated transcripts.
- Even a modest defect in NNS regulation of *PIC2* elicits severe defects in cell growth, increases cell size and intracellular stress, and prolongs the cell cycle.

## Introduction

Survival upon exposure to adverse conditions relies on rapid reprogramming of gene expression. At the beginning of a stress response, there is a genome-wide redistribution of both the transcription machinery and RNA-binding proteins (RBPs), particularly those involved in RNA decay^1^. In budding yeast, RNA polymerase II (Pol II) transcription is terminated by at least two pathways, which are contingent on the binding of distinct sets of proteins. Whereas the cleavage and polyadenylation factor (CPF) complex coordinates 3, end processing of pre-mRNAs, the Nrd1-Nab3-Sen1 (NNS) complex mediates the maturation of small nuclear (snRNA) and small nucleolar RNAs (snoRNAs) as well as the degradation of cryptic unstable transcripts (CUTs)^2–5^ originating from pervasive bidirectional transcription

In addition to targeting these canonical substrates, the NNS complex attenuates several messenger RNAs (mRNAs)^5–11^. Transcription termination by NNS starts with the Nrd1-Nab3 heterodimer binding a phosphorylated serine (Ser5P) residue in the C-terminal domain of the RNA polymerase (Figure 1A). Nrd1 and Nab3 subsequently recognise their cognate sequences in the RNA and recruit Sen1 to dismantle the polymerase elongation complex^2^. The NNS complex then recruits the Trf4-Air2-Mtr4 (TRAMP) complex to add short polyadenylation signals at the 3’ end of the released transcripts. These signals flag them for degradation by the exosome^2^. Since the NNS complex generally assembles during the synthesis of the first 100 ribonucleotides^12^, the resulting transcripts are usually not longer than ∼500 nucleotides.

**Figure 1.**
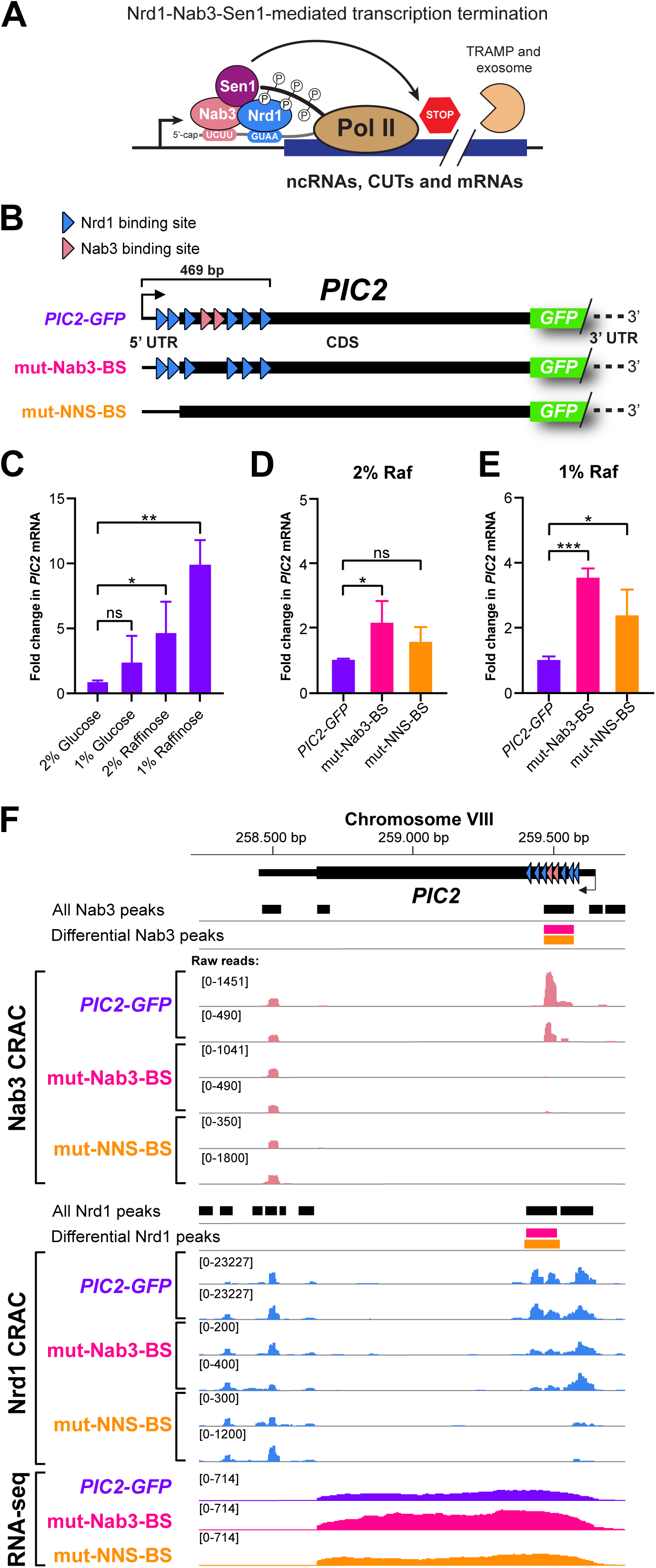
Mutating NNS-binding sites in *PIC2* increases its expression. (A) Schematic representation of NNS-mediated termination. Nrd1, Nab3 and Sen1 are co-transcriptionally recruited to RNA Polymerase II (Pol II) through an interaction with phosphorylated serine-5 in the CTD of Pol II. The binding of Nab3 and Nrd1 to specific sequences in the RNA transcript pauses Pol II and activates Sen1, which then terminates transcription. The truncated nascent transcript is targeted for degradation by the TRAMP-exosome complex. (B) Diagrammatic summary of the Nab3 and Nrd1 RNA binding sites in *PIC2* and our two modified strains: mut-Nab3-BS and mut-NNS-BS. (C) Fold changes of *PIC2* mRNA levels in the *PIC2-GFP* strain relative to 2% glucose. Bar plots display means and SDs of three independent biological repeats. *: p< 0.05, **: p< 0.01 and ***: p< 0.001 and ‘ns’: p>0.05 (unpaired t-test). (D-E) Fold changes of *PIC2* mRNA levels in the mut-Nab3-BS and mut-NNS-BS mutants, respectively, compared to the parental strain in two different raffinose concentrations. Bar plots display means and SDs of three independent biological repeats. *: p< 0.05, **: p< 0.01 and ***: p< 0.001 and ‘ns’: p>0.05 (unpaired t-test). (F) Removing Nab3 binding sites in *PIC2* does not prevent Nrd1 binding. Snapshots of read distributions obtained in Nab3 CRAC (top panel), Nrd1 CRAC (middle panel) and RNA-seq data (lower panel) of the parental *PIC2-GFP* strain and mut-Nab3-BS and mut-NNS-BS mutants. CRAC datasets show results from two independent repeats. To compensate for differences in library coverage, the raw reads shown in the y-axes were adjusted to display similar signals in neighbouring peaks that had not been deemed differentially bound by the DBPeaks package. The RNA-seq data track was generated by merging biological triplicate datasets. Nrd1 and Nab3 binding sites were identified using pyCalculateFDRs from the pyCRAC package^10^. Differential binding sites identified by DBPeaks in mut-Nab3-BS (pink) and mut-NNS-BS (orange) are indicated.

To orchestrate the gene expression changes required to endure nutrient deprivation, the NNS complex globally changes its occupancy^11,13^. Surprisingly, under these conditions, transcripts from some induced genes are simultaneously targeted by NNS^11,13^. This implies that a fraction of the mRNAs produced during stress is destined to be immediately destroyed by NNS transcription termination. Additionally, the NNS complex may influence the overall levels of certain mRNAs by terminating the transcription of some cryptic unstable transcripts (CUTs). When a CUT and an mRNA are transcribed in the same direction, non-productive transcription starting from the CUT initiation site negatively correlates with transcription initiation from the distal mRNA site^14^. Although this mode of regulation and its activation remain unconfirmed, previous studies^9,14,15^ have shown that the re-direction of the RNA polymerase II to the protein-coding transcription start site is linked to nucleotide availability.

Given the stochastic nature of Pol II transcription, we conjectured that transcribing stress-responsive genes and concurrently targeting this transcription for premature termination may have evolved as a strategy to reduce overshooting of gene expression during nutrient scarcity. We speculated that this adjustment would be particularly relevant to genes encoding proteins which, when present at high levels, would pose a disadvantage to cells facing challenging environments.

To test these hypotheses, we focused on *PIC2,* an NNS target that is strongly upregulated during glucose starvation^11^ and encodes a mitochondrial phosphate and copper importer^16,17^. Primarily, *PIC2* was selected as a model target because inadequate expression of its protein causes a clear growth defect against which we could benchmark our observations^17^. Furthermore, because Pic2 is functionally conserved in higher eukaryotes, we could test whether altered expression of *SLC25A3*, the homologue in mammals whose systemic knock-out and overexpression have been associated with disease^18,19^, caused similar phenotypes in yeast.

Here, we demonstrate that even a relatively small increase in *PIC2* mRNA levels caused by disrupting NNS regulation of PIC2 in budding yeast has significant physiological consequences. Incomplete binding of the NNS complex to *PIC2* results in decreased growth, defects in energy homeostasis, increased cell size and delayed cell cycle progression. Our phenotypic analysis suggests that the last two defects are primarily due to impaired NNS termination at other targets.

Overall, we show that a minor imbalance in the RNA recognition capabilities of Nab3 and Nrd1 for a single cognate transcript is sufficient to cause differential binding among co-regulated NNS targets. This imbalance acts as a pathophysiological mechanism that triggers cellular malfunction and reduces cellular fitness.

## Results

### *PIC2* expression is increased in mutants lacking NNS-binding sites

To quantitatively monitor levels of Pic2, we used a strain in which *PIC2* was expressed as a C-terminal GFP-fusion protein (*PIC2-GFP*). We subsequently used CRISPR/Cas9 gene editing to introduce silent point mutations in previously reported *PIC2* NNS binding sites^11^ (Figure 1B). We engineered two mutants: one without Nab3 binding sites in *PIC2* (mut-Nab3-BS) and another without both Nrd1 and Nab3 binding sites (mut-NNS-BS; Figure 1B). We expected that blocking NNS binding within the first 500 bp downstream of the *PIC2* transcription start site (Figure 1B) would prevent its premature termination during transcription and increase *PIC2* mRNA and protein levels. *PIC2*, which encodes a mitochondrial copper and phosphate transporter important for respiration^16,17^, is downregulated during fermentation^11^. Thus, we supplemented growth media with high (2%) or low (1%) concentrations of the poorly fermentable sugar raffinose. In raffinose, cells respire and, therefore, must upregulate *PIC2* expression. Consistently, RT-qPCR analyses on RNA extracted from yeast grown in raffinose or glucose confirmed that *PIC2* expression is substantially higher in raffinose (Figure 1C).

We next checked whether altering NNS binding to *PIC2* indeed increased its mRNA abundance. RT-qPCR revealed that *PIC2* mRNA levels increased by 2- to 4-fold in mut-Nab3-BS compared to its parental strain (Figures 1D-E), consistent with defective premature transcription termination of *PIC2*. Since this upregulation was highest (∼4-fold) in lower raffinose concentrations (Figure 1E), we performed subsequent experiments in 1% raffinose. Unexpectedly, *PIC2* mRNA levels only modestly increased in mut-NNS-BS (Figures 1D-E), possibly because its larger number of nucleotide substitutions affected mRNA stability, overshadowing any effects from diminished NNS regulation.

To ascertain that the mutations introduced in *PIC2* indeed impeded NNS binding, we mapped Nab3 and Nrd1’s transcriptome-wide occupation in the parental and mutant *PIC2-GFP* strains using the cross-linking and analysis of cDNAs method^20,21^ (CRAC, Figures S1A-B). To identify statistically significant changes in the NNS-binding profiles, we developed DBPeaks, a Python peak-calling script that employs pyCRAC^10^, BEDTools^22,23^ and DESeq2^24^ to detect differentially bound peaks among CLIP/CRAC datasets (Figures S1A and 1F; tracks showing differential peaks). As expected, disrupting both Nrd1 and Nab3 binding sites (mut-NNS-BS) almost completely abrogated the binding of both proteins to the *PIC2* transcript. The mut-Nab3-BS mutations blocked Nab3 binding to *PIC2*, as hypothesised, but Nrd1 cross-linking was predominantly still retained throughout its binding sites. These data demonstrate that Nrd1 can still be recruited to *PIC2* even when Nab3 no longer binds the transcript (Figure 1F).

### NNS repression of target mRNAs is broadly applicable

Although *PIC2* mRNA levels in low raffinose medium were 10-fold higher than those grown in glucose-containing medium (Figure 1C), the gene is still lowly expressed relative to other transcripts (Figure 2A). Therefore, the NNS attenuation mechanism may be limited to genes with low transcription. To address this concern, we engineered reporter constructs where the expression of *GFP* was driven from the strongly inducible galactose-responsive *GAL1* promoter. The design of our constructs (Figure 2B) was inspired by an *in vivo* screen that identified optimal sequence determinants for NNS-mediated transcription attenuation in non-coding RNAs (i.e., the ‘supermotif’)^25^. We designed a *pGAL1-GFP* construct containing the 5’UTR of *RPL20A*, a highly expressed gene (Figure 2A), in which canonical Nrd1 or Nab3 RNA-binding motifs were removed (Figure 2B; BSWT). Within the BSWT parental construct, we embedded either one (BS1) or two (BS4) supermotifs into the 5’ UTR. Additionally, we developed variants in which we swapped the first Nab3 and Nrd1 motif in the supermotif (BS3) or substituted the spacer regions of the supermotif with native sequences (BS2), changes previously shown to diminish NNS attenuation^25^.

**Figure 2.**
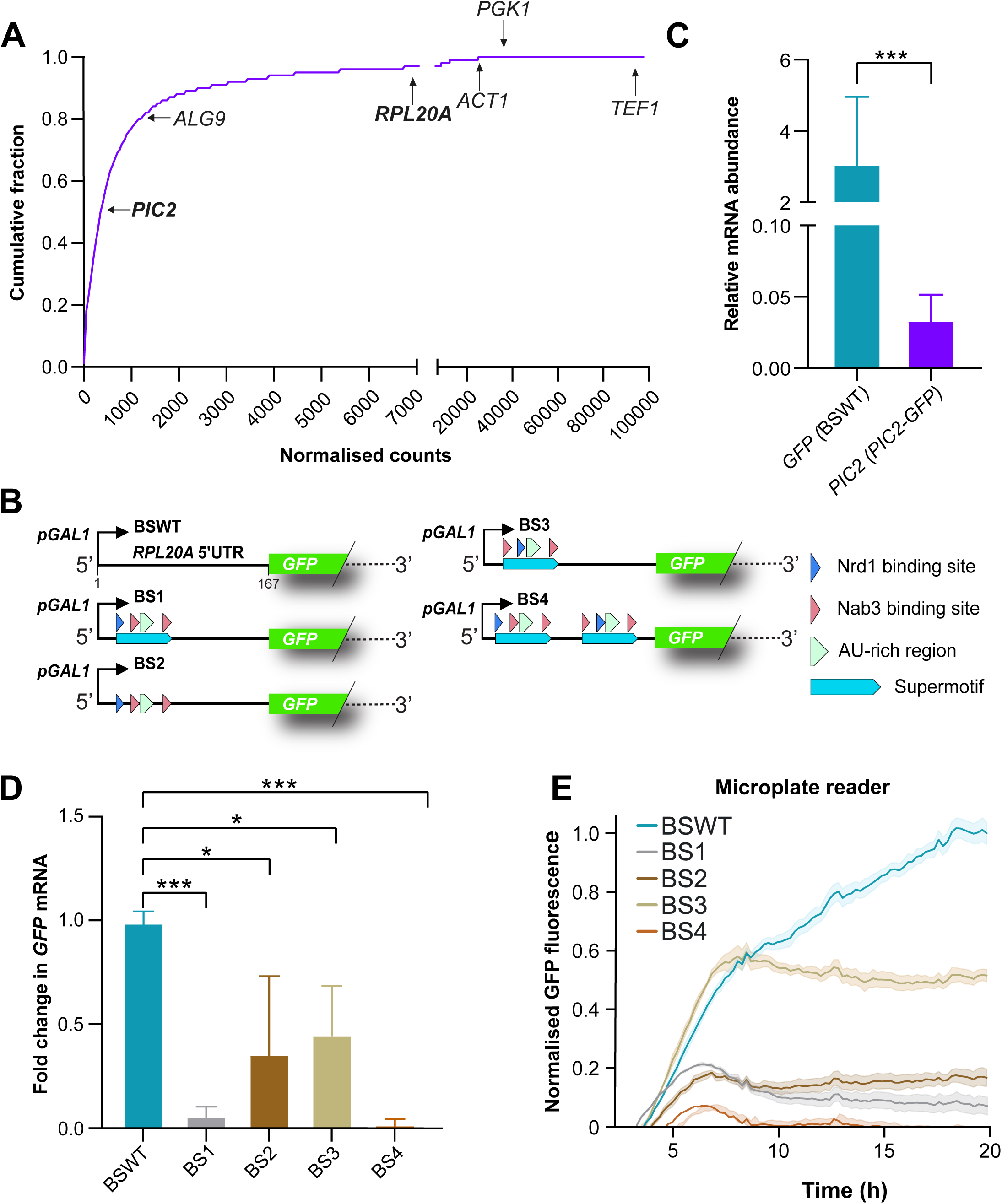
NNS attenuation of target protein-coding genes applies generally. (A) *PIC2* is a lowly expressed gene. Cumulative distribution of transcript abundance quantified by RNA-seq in the *PIC2-GFP* strain grown in low raffinose medium. Indicated in the cumulative plot are the abundances of not only *PIC2* but also the level of the reference genes used in our RT-qPCR analyses (*ALG9*, *PGK1*, *ACT1* and *TEF1*) and *RPL20A*, of which the 5’ UTR was used to generate the synthetic constructs shown in. (B) Nab3 and Nrd1 RNA binding sites in the *pGAL1-GFP* synthetic reporter constructs. In the BSWT strain, the *GFP* reporter is preceded by a *pGAL1* promoter and the 5’UTR of *RPL20A*. BS1 contains one supermotif in the 5’ UTR, BS2 encodes the same supermotif with replaced spacer sequences, BS3 codes for the supermotif with swapped NNS binding sites, and BS4 encompasses two supermotifs upstream of the coding sequence. (C) Relative mRNA abundance of *GFP* and *PIC2* in the BSWT and *PIC2-GFP* strains. Whilst BSWT cultures were grown in 1% sucrose and 1% galactose, *PIC2-GFP* ones were in 1% raffinose. ***: p< 0.001 (unpaired t-test). (D) Changes in *GFP* mRNA across the strains encoding different synthetic expression systems. Means and SDs from three biological repeats are shown. *: p< 0.05, **: p< 0.01 and ***: p< 0.001. Cells were grown in 1% sucrose and 1% galactose medium. (E) Normalised GFP fluorescence measurements in the reporter strains recorded in a microplate reader across time. Cells were grown in medium containing 1% sucrose and 1% galactose.

After integrating these constructs into the *GAL2* locus, we quantified *GFP* mRNA levels of the resulting strains using RT-qPCR (Figures 2C-D) and GFP protein abundance in a microplate reader (Figure 2E). The relative abundance of *GFP* transcripts in the BSWT strain was two orders of magnitude larger than that of *PIC2* mRNA in the *PIC2-GFP* parental strain (3.09 vs 0.03), demonstrating that this synthetic system generated highly expressed transcripts (Figure 2C). Inserting even a single supermotif in the 5’ UTR considerably lowered the mRNA and protein levels of the *GFP* reporter (Figures 2D-E). Constructs with replaced spacers (BS2) or swapped NNS binding sites (BS3) exhibited intermediate expression levels.

These experiments confirm that NNS-mediated attenuation and its associated reduction in gene expression extend beyond lowly expressed mRNAs and can affect protein-coding genes with strong promoters. They suggest that this regulatory role of the NNS complex could be widespread.

### Abrogating Nab3 binding to *PIC2* decreases fitness, enlarges cell size and delays cell cycle progression

To evaluate the phenotypical impact of disrupting *PIC2* NNS regulation, we asked whether the mutants displayed growth defects. After quantifying the maximum growth rates of the *PIC2-GFP*, mut-Nab3-BS and mut-NNS-BS strains (Figure 3A), we found that removing Nab3 binding sites in *PIC2* caused clear growth defects in low raffinose medium. Upon microscopic inspection of the mut-Nab3-BS cells (Figure 3B), we noticed that these cells were visibly larger than the parental *PIC2-GFP* strain. This prompted us to compare the cell sizes of the strains using flow cytometry. Indeed, the average cell size of mut-Nab3-BS cells was around 50% larger than the parental strain, but mut-NNS-BS did not show significant size increases (Figures 3C-D). To substantiate these results, we applied single-cell microfluidics and time-lapse microscopy to quantify cell size and Pic2-GFP expression levels in the parental and mutant strains^26,27^. As a negative control, we performed the same experiment with the non-fluorescent parental strain, which was used to correct for autofluorescence. Consistent with our flow cytometry data, our microfluidics analyses showed that the mut-Nab3-BS strain has a markedly larger cell volume compared to the parental strain (Figure 3E). Moreover, the resulting quantification of Pic2-GFP fluorescence in each cell revealed that the mean fluorescence, an estimate of concentration, increased in both mutant strains (Figure 3F). We conclude that NNS regulation of *PIC2* is important for controlling both cell volume and Pic2 protein concentration.

**Figure 3.**
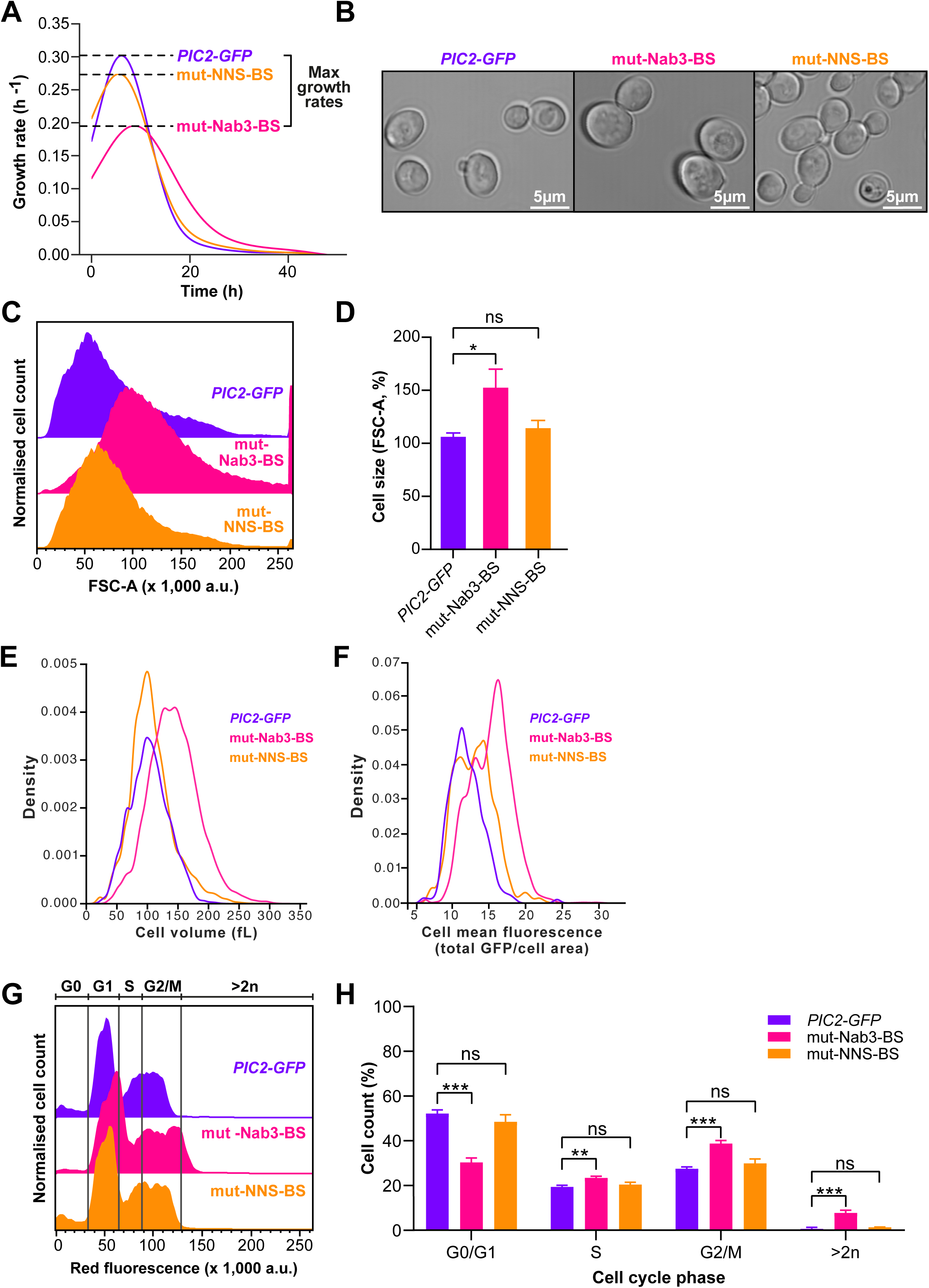
Preventing Nab3 binding to *PIC2* causes growth defects, increases cell size, enhances Pic2-GFP expression and delays cell cycle progression. (A) Time-resolved growth rates of the *PIC2-GFP*, mut-Nab3-BS and mut-NNS-BS strains. We show the mean of three technical replicates. Maximum growth rates for each strain are indicated. (B) Confocal bright-field images of *PIC2-GFP*, mut-Nab3-BS and mut-NNS-BS strains. mut-Nab3-BS are visibly larger. (C) Representative traces of the cell size distribution of *PIC2-GFP* and derived mutants from flow cytometry analyses. The sizes were compared using their forward light scatter (FSC-A) values. mut-Nab3-BS cells are larger. (D) Weighted means and SDs of the median FSC-A values for three independent biological repeats calculated for each strain and normalised to the parental *PIC2-GFP* strain. (E) Distribution of cell volumes (in fL) for the *PIC2-GFP*, mut-Nab3-BS and mut-NNS-BS strains, calculated using the microfluidics imaging data. (F) Distribution of mean cell Pic2-GFP fluorescence, calculated by dividing the Pic2 fluorescence (arbitrary units) of each cell by its area. Accounting for the larger size of the mut-Nab3-BS mutant, the concentration of GFP is still higher. (G) Representative flow cytometry traces of cells with fluorescently dyed DNA of *PIC2-GFP*, mut-Nab3-BS and mut-NNS-BS strains at mid-log phase (OD_600_ ∼0.5). (H) Weighted means and SDs of the population fraction of each strain in (E) within each cell cycle phase. Values based on biological triplicates. *: p< 0.05, **: p< 0.01 and ***: p< 0.001 and ‘ns’: p>0.05 (unpaired t-test). All shades represent half the interquartile range.

Before further exploring this phenotype, we sought to ensure that the observed cell size increase was indeed genetically linked to the mutations we introduced to disrupt NNS regulation of *PIC2* and not due to any CRISPR-related off-target effects. To this end, fluorescent strains were backcrossed with a non-fluorescent strain of the opposite mating type (BY4742). After undergoing meiosis and sporulation, the resulting diploid strain yielded tetrads enclosing four haploid daughter cells (Figure S2A). Among these, two encoded a wild-type version of *PIC2,* and the remaining ones expressed the *GFP*-tagged *PIC2*. Upon tetrad dissection, cell size measurements confirmed that haploids carrying the mut-Nab3-BS mutations were also larger than their non-fluorescent counterparts. Accordingly, we conclude that the increased cell size co-segregated with the disruption of NNS regulation of *PIC2*.

To substantiate this control experiment, we studied additional independently derived mutant clones. After performing RT-qPCR analyses and cell size measurements on a second mut-Nab3-BS clone (Nab3 BS #2; Figure S2B), we observed comparable increases in *PIC2* mRNA levels (Figure S2C) and cell size (Figures S2D-E). When screening for additional mut-NNS-BS clones, we serendipitously identified a strain that lacked all Nab3 binding sites and half of the Nrd1 motifs (Figure S2B; partial mut-NNS-BS). This mutant exhibited intermediate increases in *PIC2* mRNA levels (Figure S2C) and cell size (Figure S2D-E). Hence, these findings suggest that the extent of *PIC2* upregulation and the accompanying cell size enlargement was (at least in part) dictated by the number of intact Nrd1 motifs present in *PIC2,* together with the absence of Nab3 binding sites.

Earlier work has established that cell cycle progression is prolonged in oversized cells^28^. Therefore, we analysed the cell cycles of the parental and mutant *PIC2-GFP* strains using flow cytometry and propidium iodide dye to stain DNA. Notably, a greater portion of the mut-Nab3-BS populations were in S or G2/M phases (Figures 3G-H and S2F-G). Thus, our results revealed a mild cell cycle prolongation in strains lacking *PIC2* Nab3 binding sites. Moreover, a modest increase in the fraction of polyploid cells was also observed in mut Nab3 strains, suggesting a higher rate of mitotic defects. In line with our cell size measurements, the partial NNS mutant (partial mut-NNS-BS) showed intermediate cell cycle delays, and no defects were detectable in the strain lacking all NNS binding sites in *PIC2* (mut-NNS-BS; Figures 3G-H and S2F-G).

Collectively, these data show that the observed increases in *PIC2* transcript and protein levels, caused by specifically disrupting Nab3 binding to *PIC2*, result in abnormal cell size and delays in cell cycle progression.

### Impeding Nab3 binding to *PIC2* causes mitochondrial hyperpolarisation and resistance to oxidative stress

We next sought to elucidate whether the growth defects of mut-Nab3-BS in low raffinose medium were linked to the increased expression and activity of the mitochondrial copper uniporter and phosphate symporter Pic2. We first measured mitochondrial membrane potential (Δψ*_m_*), a major index of mitochondrial energy homeostasis^29^. Live cells were equilibrated with tetramethyl rhodamine ethyl ester (TMRM), a dye that accumulates in active mitochondria proportionally to their activity. Subsequent live cell confocal microscopy revealed mitochondrial membrane hyperpolarisation in the mut-Nab3-BS mutant (Figure 4A). Increased mitochondrial membrane potential can stem from enhanced respiratory rate or inhibited ATP synthesis, but the latter would cause a decreased respiration rate. Hence, to further interpret these findings, we quantified oxygen consumption. This revealed that respiratory rates were significantly increased in mut-Nab3-BS compared to the parental *PIC2-GFP* in low raffinose concentrations (Figure 4B), suggesting that the higher mitochondrial membrane potential is driven by an increased respiratory rate.

**Figure 4.**
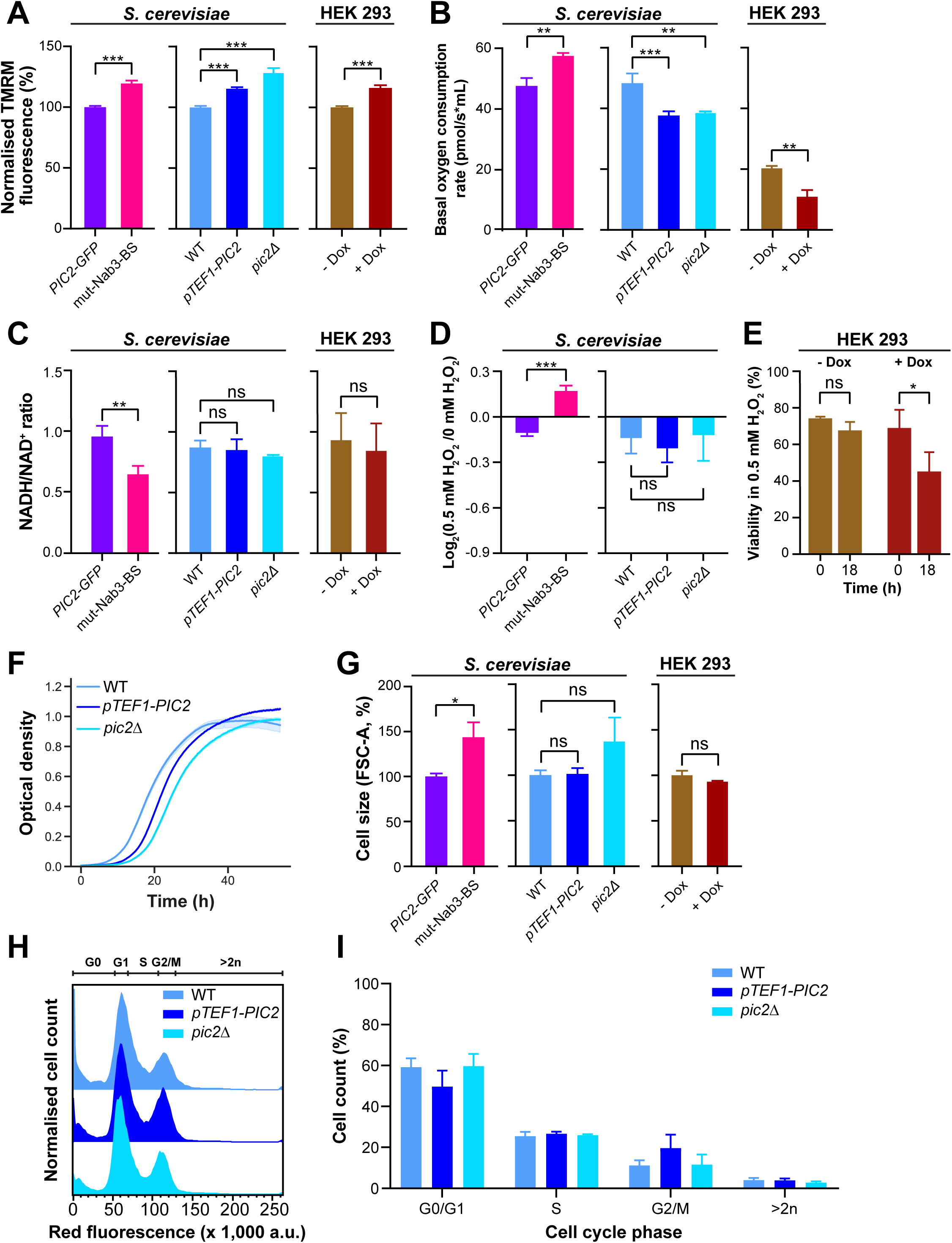
Overexpression of Pic2 decreases yeast and mammalian cell fitness but affects neither cell size nor cell cycle. (A) Mean and SEM for TMRM fluorescence values, indicating mitochondrial membrane potential, gathered from confocal imaging of more than 50 cells from three different biological repeats. (B) Mean and SD for basal oxygen consumption rates measured in biological triplicates. (C) Mean and SD for NADH/NAD^+^ ratios recorded in three independent assays. (D-E) Mean and SD of the log2-transformed fold change in the growth rate of cells grown with and without the presence of 10 mM hydrogen peroxide in the medium. Measurements were performed in a microplate reader. For HEK 293 cells (E), the mean and SD of live cells are shown. Trypan blue assays were performed in three independent samples. (F) Time-resolved optical density measurements depicting the mean and with shading half the interquartile range of three technical repeats. WT displayed the shortest lag phase. (G) Weighted means and SDs of the cell size (FSC-A values in flow cytometry) for three independent biological repeats were calculated for each cell type and normalised to that of the pertinent reference. (H) Representative flow cytometry traces of the DNA profile of *pTEF1-PIC2*, *pic2*Δ and their parental strain (WT). (I) Weighted means and SDs of the population fraction of each strain in (H) undergoing each cell cycle phase. *: p< 0.05, **: p< 0.01 and ***: p< 0.001 and ‘ns’: p>0.05 (unpaired t-test).

Based on these data and on previous work reporting that oversized cells undergo continual stress responses^28^, we hypothesised that the observed defects in energy homeostasis arose from intracellular stress. To test this, we quantified the nicotinamide adenine dinucleotide (NAD) pools of both strains by performing UV fluorescence assays in a microplate reader. Consistent with increased respiration and enhanced ROS production, NAD species in mut-Nab3-BS were predominantly in their oxidised NAD^+^ state compared to the parental strain (Figure 4C). Furthermore, to corroborate that mut-Nab3-BS was inherently stressed, we devised an oxidative stress assay based on acquired stress resistance: since exposure to a mild insult confers some resistance to subsequent ones^30^, we expected stressed cells to be less affected by environmental challenges. Indeed, whereas the growth of the parental strain was reduced by exposure to hydrogen peroxide, that of mut-Nab3-BS did not decrease (Figure 4D).

We conclude that disrupting Nab3 binding to *PIC2* mRNA causes energy homeostasis anomalies that generate greater respiratory demands and intracellular oxidative stress.

### Pic2 overexpression decreases fitness but does not alter cell size or cell cycle progression

We then aimed to identify the mechanism underpinning the described growth, cell size, cell cycle and energy homeostasis defects. The simplest justification for these anomalies was that they merely emerged from an increase in the abundance and activity of the Pic2 protein. If higher expression of Pic2 was indeed driving the described phenotypes, a strain overexpressing Pic2 in low raffinose medium should exacerbate (or at least recapitulate) all defects.

To span the scope of phenotypic consequences caused by changes in *PIC2* expression, we generated an overexpression mutant (*pTEF1-PIC2*; Figure S3A) and a knock-out strain (*pic2*Δ; Figure S3A). In agreement with previous work^17^, the *pic2*Δ mutant suffered a growth defect in low raffinose medium, with cells taking longer to enter their exponential growth phase (Figure 4F). Similarly, overexpressing *PIC2* caused a comparable growth delay (Figure 4F). We confirmed the incurred fitness costs for these mutants with competition assays, which showed that cells lacking or overexpressing *PIC2* were eventually outnumbered by their parental counterparts (Figure S3B).

We then measured the mitochondrial membrane potential and oxygen consumption of these strains. Like mut-Nab3-BS, the *PIC2* knock-out and overexpressing strains displayed mitochondrial hyperpolarisation (Figure 4A). However, respirometry measurements and NAD pool evaluations established that the origin of this mitochondrial hyperpolarisation was different to that affecting mut-Nab3-BS: basal oxygen consumption of *pic2*Δ and *pTEF1-PIC2* was reduced (Figure 4B), and neither significant NADH/NAD^+^ changes nor oxidative stress resistance were detected in any of these mutants (Figures 4C-D). Increased mitochondrial membrane potential can only co-exist with decreased respiratory rate if ATP synthesis is inhibited. In turn, compromised ATP generation could potentially be caused by altered phosphate homeostasis in our *PIC2* overexpression model.

In summary, these findings verify that even subtle deviations from optimal levels of Pic2 expression can inhibit respiration, thereby diminishing cell fitness.

### Overexpression of human *PIC2* causes energy homeostasis defects

Throughout this work, we focused on *PIC2* partially because mutations in the human Pic2 orthologue, *SLC25A3*^31^, are linked to cardiac and muscular diseases^32^ and, like *PIC2*, the expression of *SLC25A3* is tightly controlled^33^. Despite not possessing Nrd1 and Nab3 orthologues, mammalian cells employ microRNAs to fine-tune gene expression^34^. Indeed, in humans, miR-144 is a key regulator of *SLC25A3*^35^. Similarly to *pic2*Δ yeast, mammalian cell models lacking *SLC25A3* show mitochondrial hyperpolarisation^36^ and decreased ATP synthesis^18^. Thus, we investigated whether overexpressing *SLC25A3* would lead to changes in mitochondrial activity resembling that of our *PIC2-*overexpressing yeast model. We used the Flp-In™ system to generate a HEK 293 cell line in which overexpression of *SLC25A3* could be induced with doxycycline (Figures S3C-E). As expected, the mammalian *SLC25A3*-overexpressing model fully recapitulated the mitochondrial hyperpolarisation (Figure 4A) and reduced respiration (Figure 4B) phenotypes observed in yeast. Additionally, cells overexpressing *SLC25A3* maintained a NADH/NAD^+^ status similar to that of their uninduced counterparts (Figure 4C) and showed reduced viability under oxidative challenges (Figure 4E).

We conclude that suboptimal Pic2 levels in yeast and human cells cause energy homeostasis defects, hindering respiration and causing decreased fitness. Importantly, however, altered expression of *PIC2* did not cause significant changes in yeast or human cell size (Figure 4G) nor significantly affected yeast cell cycle progression (Figures 4H-I). Therefore, these results imply that the observed cell size abnormalities in mut-Nab3-BS are not directly linked to increased Pic2 protein levels.

### Blocking premature NNS termination of *PIC2* increases cell size and delays cell cycle progression

Previous studies^15,37^ reported cell size increases in mutants in which Nab3 or Nrd1 were prevented from binding their RNA targets. Accordingly, we hypothesised that the cell size abnormalities in mut-Nab3-BS could be linked to changes in the availability of Nrd1 and Nab3. To test this possibility, we employed a strain in which Nab3 could be conditionally depleted from the nucleus in the presence of rapamycin^11,38^ (*NAB3-FRB*; Figure 5A) with the parental strain (WT) as a negative control (Figure 5B). Using flow cytometry, we monitored cell size at 1, 4, and 8 hours after adding rapamycin. Cell size steadily increased after 4 hours of rapamycin treatment, and after 8 hours of exposure to the drug, it became equivalent to that observed in mut-Nab3-BS. Treating the parental strain with rapamycin for 8 hours resulted in minor increases in cell size (Figures 5A and S4A). Rapamycin can arrest cells in G1^39^, ultimately increasing their cell size^28^. Thus, although engineered to resist the toxicity of rapamycin^38^, the parental strain does not appear to be completely insensitive to the drug.

**Figure 5.**
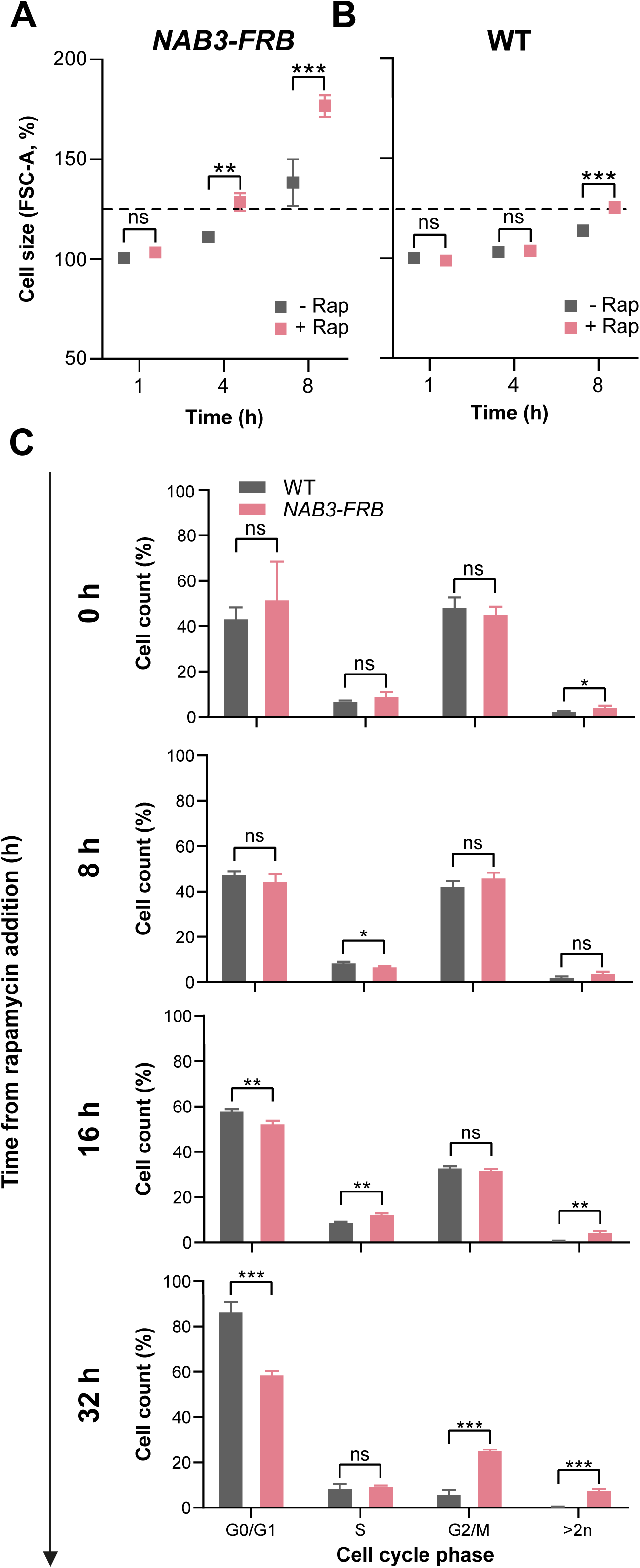
Depleting Nab3 from the nucleus increases cell size and prolongs the cell cycle. (A-B) Time-resolved means and SDs of cell size medians from three independent evaluations of *NAB3-FRB* and its parental strain (WT). The resulting values were normalised to that of the untreated reference collected in the first time point. (C) Bar plots displaying weighted means and SDs of the population fraction of *NAB3-FRB* and its WT control within each cell cycle phase. *: p< 0.05, **: p< 0.01 and ***: p< 0.001 and ‘ns’: p>0.05 (unpaired t-test).

To test whether, in addition to cell size, the cell cycle was also disrupted upon nuclear depletion of Nab3, we performed time-resolved cell cycle analyses in the Nab3-depleted mutant and its parental strain (Figures 5C and S4B). Since yeast doubles every 3.5 hours in low raffinose medium, we extended the timespan of the experiment to encompass several cell cycles through which the oversized mutant was expected to become delayed. Indeed, 16 hours after sequestering Nab3 in the cytoplasm, cells spent less time in G1, underwent prolonged S phases and became polyploid more frequently than their parental counterparts (Figures 5C and S4B). These effects aggravated after 32 hours, when significantly longer S phases were replaced by extended G2/M phases, reminiscent of cell cycle changes in the mut-Nab3-BS strain. As anticipated, rapamycin eventually also arrested the parental strain in G1^39^, thereby gradually shifting its population towards 1n states throughout the experiment.

These data show that nuclear depletion of Nab3 results in cell size increases and aberrant cell cycle progression. Surprisingly, these phenotypes echo the phenotypes observed in the mut-Nab3-BS strain, which contains only two single nucleotide substitutions that disrupt Nab3 binding to one mRNA.

### Disrupting Nab3 binding to *PIC2* mRNA alters Nrd1 binding to NNS targets

Based on these observations, we propose a ‘sequestration model’ to explain the observed cell size and cell cycle defects in the mut-Nab3-BS strain. Although mutations in mut-Nab3-BS prevented Nab3 binding to *PIC2* mRNA (Figure 1C), Nrd1 still binds the *PIC2* transcript (Figure 1C) and might recruit Sen1. We hypothesised that because the *PIC2* transcripts in mut-Nab3-BS could presumably no longer be terminated, Nrd1 could be sequestered to overabundant *PIC2* mRNAs by either titration or prolonged retention to the transcripts (see Discussion). Either scenario would reduce the availability of Nrd1, thereby altering its binding across its usual targets. This hypothesis was based on previous work^40^ demonstrating two key points: (1) RNA degradation is required for the effective release of Nrd1 and Nab3 from their substrates, and (2) overexpression of a ncRNA decoy that cannot be degraded was shown to disrupt NNS termination across the entire transcriptome.

To explore this sequestration model, we conducted RNA-seq and proteomics analyses on *PIC2-GFP* and mut-Nab3-BS (Table 1). Our findings revealed that 488 transcripts (4% of all detected RNAs) and 92 proteins (9% of all identified factors) were differentially expressed in mut-Nab3-BS (Figures 6A-B). In contrast, mut-NNS-BS, which did not present strong phenotypes, showed changes in only 41 transcripts and 50 proteins (0.1% and 4.7% of totals; Table 1). Importantly, none of the NNS components were differentially expressed in our transcriptomic and proteomic datasets, suggesting that the observed changes in gene expression are not due to alterations in NNS protein levels.

**Figure 6.**
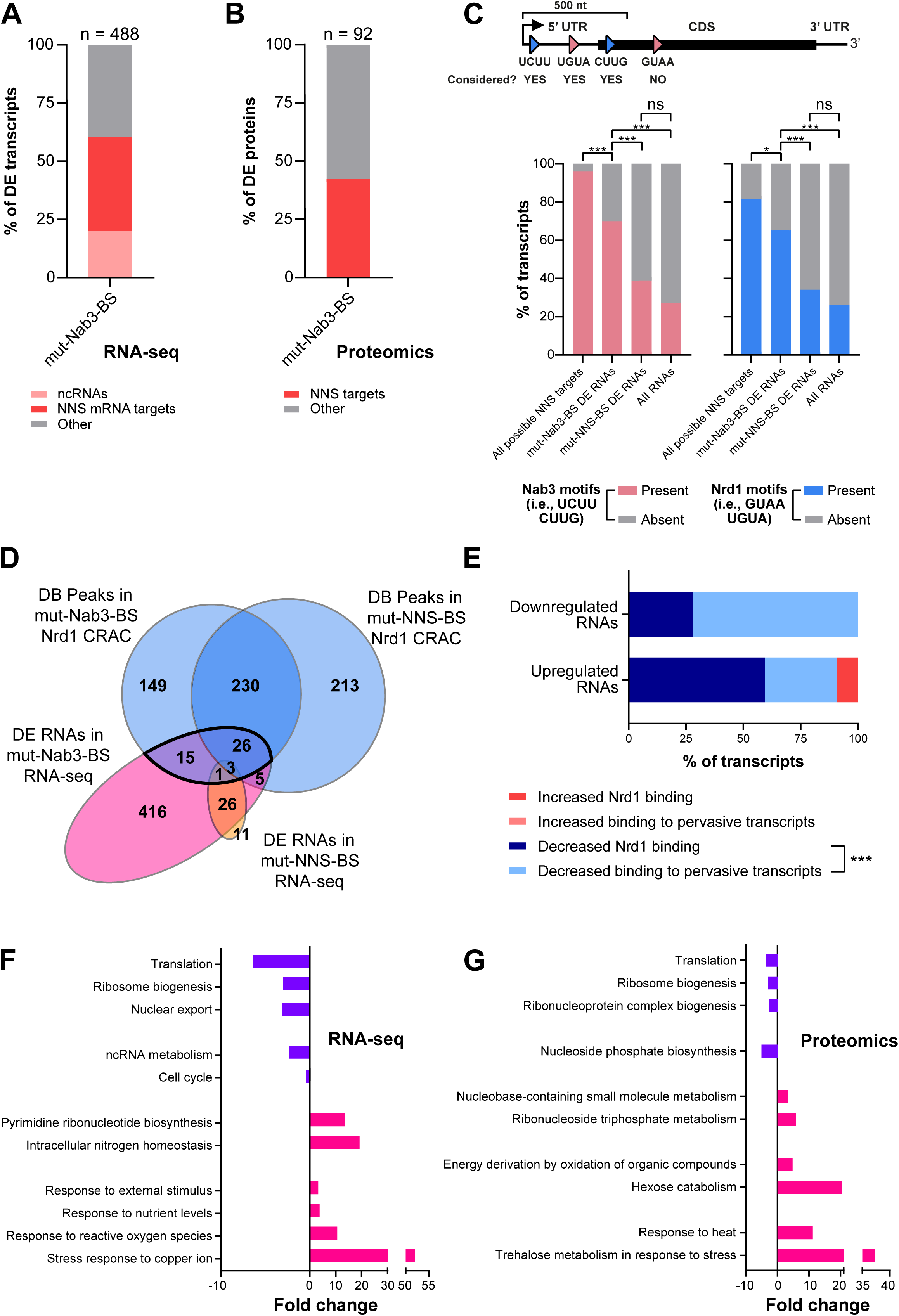
Disrupting Nab3 binding to *PIC2* prompts an accumulation of NNS targets at a transcriptomic and proteomic level. (A-B) Stacked bar charts showing the proportion of differentially expressed RNAs and proteins in mut-Nab3-BS bound by Nab3 and Nrd1 in our CRAC data. (A) also indicates the portion of differentially expressed transcripts comprising ncRNAs. (C) Fraction of transcripts containing Nab3 and Nrd1 binding motifs across differentially expressed transcripts of the mut-Nab3-BS and mut-NNS-BS strains, all transcripts annotated in the genome (negative control, ‘All RNAs’) and all transcripts with at least one Nrd1 or Nab3 motif (positive control, ‘All possible NNS targets’). As shown in the schematic, the analysis only considered Nrd1 or Nab3 motifs within the first 500 nucleotides downstream of the gene start site. (D) Venn diagrams showing overlaps between the differentially expressed (DE) transcripts and the differentially bound (DB) Nrd1 peaks in mut-Nab3-BS and mut-NNS-BS. RNAs that are differentially expressed and differentially bound by Nrd1 in mut-Nab3-BS are itemised in (E). (E) Stacked bar charts displaying the nature of the change of the differentially bound Nrd1 peaks (increased or decreased) across differentially expressed (DE) RNAs (upregulated or downregulated) in the mut-Nab3-BS strain. DB peaks lay directly within the first 500 nucleotides of the DE RNA or were detected within a pervasive transcript expressed just upstream of the DE RNA). (F-G) Gene ontology enrichment analyses among differentially expressed transcripts and proteins in mut-Nab3-BS. Only terms with FDR < 0.05 were considered. *: p< 0.05 and ***: p< 0.001 (Fisher’s exact test).

**Table 1.**
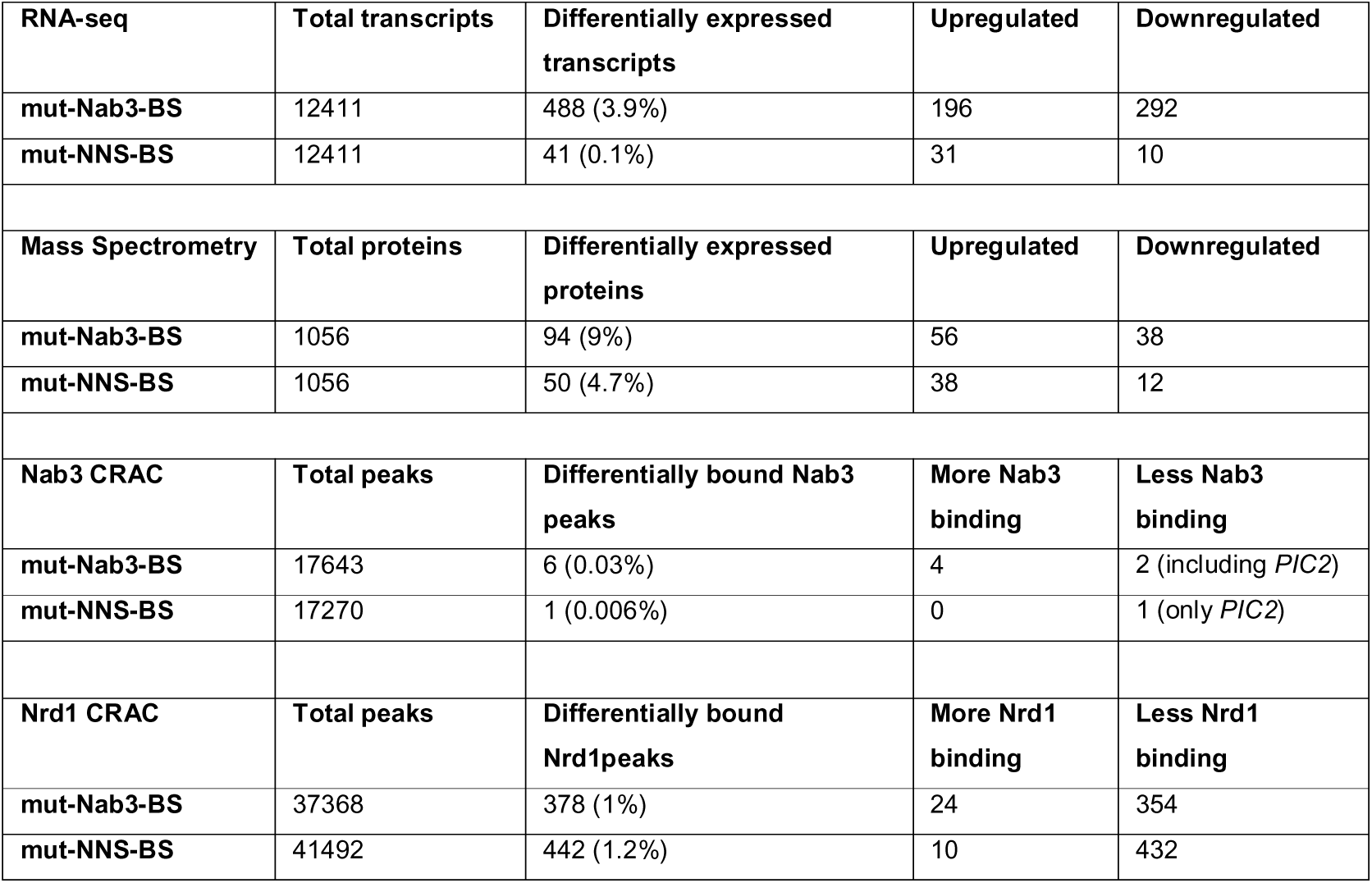
Summary of the key findings of the RNA-seq, mass spectrometry and CRAC experiments.

| RNA-seq | Total transcripts | Differentially expressed transcripts | Upregulated | Downregulated |
| --- | --- | --- | --- | --- |
| mut-Nab3-BS | 12411 | 488 (3.9%) | 196 | 292 |
| mut-NNS-BS | 12411 | 41 (0.1%) | 31 | 10 |
| Mass Spectrometry | Total proteins | Differentially expressed proteins | Upregulated | Downregulated |
| mut-Nab3-BS | 1056 | 94 (9%) | 56 | 38 |
| mut-NNS-BS | 1056 | 50 (4.7%) | 38 | 12 |
| Nab3 CRAC | Total peaks | Differentially bound Nab3 peaks | More Nab3 binding | Less Nab3 binding |
| mut-Nab3-BS | 17643 | 6 (0.03%) | 4 | 2 (including *PIC2*) |
| mut-NNS-BS | 17270 | 1 (0.006%) | 0 | 1 (only *PIC2*) |
| Nrd1 CRAC | Total peaks | Differentially bound Nrd1 peaks | More Nrd1 binding | Less Nrd1 binding |
| mut-Nab3-BS | 37368 | 378 (1%) | 24 | 354 |
| mut-NNS-BS | 41492 | 442 (1.2%) | 10 | 432 |

If our Nrd1 sequestration model is correct, then changes in Nrd1 availability should also impact the expression of other NNS targets. Accordingly, we hypothesised that many of the differentially expressed transcripts in the mutant lacking Nab3 binding sites in *PIC2* (mut-Nab3-BS) would be regulated by the NNS complex. Supporting this hypothesis, over 60% of the differentially expressed transcripts in mut-Nab3-BS were also found to be bound by Nrd1 and Nab3 in our CRAC data (Figure 6A). Of these transcripts, approximately 20% were non-coding RNAs, including numerous cryptic unstable transcripts (CUTs), while around 40% were protein-coding. Similarly, more than 40% of the differentially expressed proteins in mut-Nab3-BS were identified as putative NNS targets in our CRAC datasets.

To further substantiate these findings, we asked whether the differentially expressed transcripts in the mut-Nab3-BS strain were enriched for Nrd1 and Nab3 RNA-binding motifs (Figure 6C). Indeed, these motifs were significantly more prevalent among the differentially expressed transcripts in mut-Nab3-BS compared to those in the entire transcriptome. In contrast, the differentially expressed RNAs in the mutant without NNS binding sites in *PIC2* (mut-NNS-BS) strain were not significantly enriched for Nrd1 and Nab3 motifs.

Collectively, these findings suggest that disrupting Nab3 binding to *PIC2* not only affects this specific transcript but also influences the abundance of other NNS target transcripts, indicating a broader co-regulation of NNS RNA targets across the transcriptome.

To confirm that the observed changes in the transcriptome and proteome were indeed driven by altered Nab3 and Nrd1 binding to NNS targets, we employed our DBPeaks tool to identify differentially bound Nab3 and Nrd1 peaks within the CRAC datasets (Table 1). Despite preventing Nab3 binding to *PIC2* in both mutants, mutations in the Nab3 motifs of *PIC2* did not significantly affect global Nab3 binding (Table 1). In contrast, Nrd1 binding was significantly altered in both mutants, with hundreds of differentially bound Nrd1 peaks identified (representing 1% of all detected peaks; Figure 6D; Table 1). Consistent with reduced Nrd1 availability, most of the 378 differentially bound Nrd1 sites in mut-Nab3-BS exhibited less Nrd1 binding (Table 1). Importantly, more than 10% of these sites (41) were in differentially expressed genes (Figure 6D; Table 1), such as *DUR3,* a stress-specific mRNA regulated by the NNS complex^11^ (Figure S5). As in *DUR3,* reduced Nrd1 binding predominantly occurred in regions annotated to contain pervasive transcripts (e.g., CUTs) upstream of protein-coding genes (Figure 6E). However, over 60% of the upregulated RNAs of mut-Nab3-BS exhibited decreased Nrd1 binding to their coding sequence (Figure 6E), suggesting that their abundance increases due to defective NNS termination.

While mut-NNS-BS also exhibited numerous differentially bound Nrd1 peaks, these changes did not impact the transcriptome and proteome as much as compared to mut-Nab3-BS, as evidenced by the fewer differentially expressed genes (Figure 6D; Table 1). Additionally, its differentially expressed transcripts were not significantly enriched for Nrd1 and Nab3 motifs (Figure 6C), suggesting that their altered expression is not linked to defective NNS regulation. Therefore, although both mutants demonstrated altered Nrd1 binding, the resulting transcriptomic and phenotypic consequences were markedly different.

To support these findings, we conducted gene ontology analyses on the differentially expressed transcripts and proteins of mut-Nab3-BS (Figures 6F and 6G). Consistent with the observed excessive cell size of the mut-Nab3-BS mutant, both translation and transcription processes were downregulated^28^ (Figures 6F-G). The downregulation of transcripts involved in cell cycle regulation may explain the delayed transition of mut-Nab3-BS through the G2/M phase (Figure 6F). Moreover, the upregulation of responses to nutrient, oxidative, and copper stress suggests higher levels of intracellular stress in mut-Nab3-BS and, accordingly, increased tolerance to extracellular ones (Figures 6F-G). Consistent with increased respiratory rates, energy derivation processes were also significantly enhanced (Figures 6F-G). Conversely, the same analyses in mut-NNS-BS yielded no significant enrichments for differentially expressed transcripts and among its proteome, only sugar metabolism was upregulated.

Notably, transcripts and proteins involved in ribonucleotide homeostasis were strongly upregulated in mut-Nab3-BS (Figures 6F-G). Although this biochemical route is functionally unrelated to Pic2 in *Saccharomyces cerevisiae*, several of the enzymes involved are regulated by the NNS complex^14,15^. Subsequent metabolomic analyses revealed a systemic upregulation of metabolites connected to pyrimidine synthesis in mut-Nab3-BS (Figures S6A-B). These findings indicate that the gene expression changes observed in this strain significantly altered its metabolic flux.

Cumulatively, these results demonstrate that inhibiting Nab3 binding to a single mRNA species changes Nrd1 occupancy across a small fraction of its targets (∼1%). However, this was sufficient to cause severe phenotypic consequences, including increased cell size, reduced fitness, and defects in energy homeostasis.

## Discussion

Here we propose a role for the *Saccharomyces cerevisiae* Nrd1-Nab3-Sen1 (NNS) complex as a co-transcriptional regulator for protein-coding genes that require tight control. Using the copper/phosphate carrier *PIC2* as a model NNS target, we show that Nab3 binding reduces *PIC2* mRNA abundance and that this mechanism regulating *PIC2* expression is critical for cellular fitness during stress. Furthermore, our results suggest that tight regulation of this mitochondrial transporter is evolutionarily conserved in yeast and human cells. Combining phenotypic dissection, multi-omics profiling, and transcriptome-wide NNS-RNA binding analyses, we demonstrate that disrupting the binding of Nab3 to *PIC2* causes a redistribution of Nrd1, alters the termination of co-regulated targets and leads to aberrant cell size and cell cycle progression in yeast.

These findings show that preventing a single RNA-binding protein from binding to one mRNA target can generate transcriptomic and phenotypic changes that significantly undermine a cell’s ability to withstand harsh environments.

### The Goldilocks’ modulation of PIC2 expression is conserved across eukaryotes

Our results indicate that cells displaying high Pic2 protein levels are penalised when subjected to nutrient restriction or detrimental conditions demanding efficient respiration for survival. We were able to show this in both yeast and human cells. Interestingly, respiration inhibition and increased susceptibility to extracellular ROS were also observed in mutants lacking *PIC2*. Despite also showing growth defects in our partially respirable raffinose-containing medium, *pic2Δ* cells are known to perform nearly identically to the parental strain when grown in rich media containing fermentable sugars^17^. These observations align with general microbial adaptive behaviour: while bet hedging enables the development of stochastic phenotypes that may be advantageous in ideal growth conditions and potentially beneficial in unpredictable scenarios, adaptation also requires that the expression pattern of some genes becomes ‘just right’. Thus, maintaining *PIC2* expression at a ‘Goldilocks’ level is crucial for optimal cellular function under varying environmental stresses.

Although the expression of *PIC2* and its human homologue (*SLC25A3*) is regulated to control their levels, the regulatory mechanisms involved are markedly different. In human cells, *SLC25A3* regulation involves the microRNA machinery^35^, a system that is absent in the yeast *Saccharomyces cerevisiae*. This distinction underscores a fundamental divergence in the molecular strategies evolved by these organisms to achieve gene expression homeostasis.

### How does a single change in RNA-NNS interactions affect RNA termination across many NNS targets?

We demonstrated that a 3-4-fold increase in *PIC2* mRNA levels, which is typically a lowly expressed transcript, in the mutant mut-Nab3-BS strain was sufficient to alter Nrd1 occupancy on other NNS target transcripts. This alteration led to significant phenotypic disadvantages. Although the precise molecular cascade that results in the diminished availability of Nrd1 for its target transcripts remains unclear, we hypothesise that Nrd1 titration may occur through various mechanisms:

Nab3, which possesses prion-like domains^41^, partitions into nuclear granules during nutrient starvation^42^. Despite also containing intrinsically disordered regions, Nrd1 has not been reported to engage in phase separation. Nevertheless, previous work has noted the presence of nucleoids in strains with mutations in the RNA-recognition motif of Nrd1^37^, suggesting that Nrd1 might form condensates in response to environmental stimuli, possibly in conjunction with target transcripts and Nab3. Additionally, Nrd1 and/or Sen1 may be sequestered to different subcellular compartments while still bound to accumulating *PIC2* transcripts in the mut-Nab3-BS mutant. If excess *PIC2* transcripts were translated, Nrd1 and Sen1 could be exported alongside the transcript to the cytoplasm, effectively titrating these proteins away from the nucleus. Indeed, although Nab3, Nrd1 and Sen1 localisation is predominantly nuclear, they all have been detected in ribonucleoproteins, which exhibit dynamic cellular localisation and could potentially shuttle when bound to their targets^43^.

Since Nrd1 is an abundant protein – around 19,600 molecules per cell^44^ – it seems improbable that a mild *PIC2* overabundance would significantly alter Nrd1’s ability to bind other target transcripts. Conversely, Sen1 appears to be a limiting factor for the assembly of the complex (125 molecules per cell^44^) and so tightly modulating its concentration is required for efficient NNS-mediated termination^45^. Accordingly, we posited that the most likely cause underlying disrupted NNS-termination in our system is the sequestration of Sen1 to mut-Nab3-BS *PIC2* transcripts through an interaction with Nrd1. To test this, we performed several Sen1 CRAC experiments in *PIC2-GFP* and derived mutants. Nonetheless, under the growth conditions used, Sen1 cross-linking to RNA was inefficient relative to the other NNS components (Figure S1B). Despite multiple attempts, the resulting Sen1 CRAC cDNA libraries were of too low complexity (see Data availability). This prevented us from drawing sensible conclusions about Sen1 recruitment to mRNA substrates in our mutant strains. Therefore, we were unable to confirm whether Sen1 occupancy in NNS targets was also significantly affected in mut-Nab3-BS.

### NNS regulation is integral to achieving optimal cell size and normal cell cycle progression

Our results show that impairing NNS termination of only a single mRNA can cause severe cell size anomalies. These findings further bolster a Sen1 sequestration model as restrictions in its physiological abundance are known to cause cell cycle^45^ and cell division defects^46^. Although no aberrant cell size was documented in these studies^45,46^, a mere increase in the DNA resulting from a delayed exit from the G2 phase could theoretically drive a cell size increase^28^. Notably, our mutant was not only oversized but also displayed reduced growth and fitness as well as increased intracellular stress, which coincided with the previous characterisation of other oversized models^28^. On this basis, we conclude that our cells are likely exceeding their critical DNA:cytoplasm ratio and undergoing cytoplasmic dilution, which was defined as a senescence trait^28^. Consistent with this notion, recent work uncovered that the transcriptome is more heavily regulated by the NNS complex during quiescence^47^. Given that NNS targets are predominantly non-coding, it is possible that these operate as regulators of cell size and, by extension, affect the cell cycle. However, the function of many ncRNAs in yeast remains uncharacterised.

Our work suggests that NNS-mediated attenuation of target mRNAs could be acting as a mechanism to coordinate the expression of functionally related protein-coding genes, possibly involved in cell cycle regulation, in response to environmental cues. Our results imply that the levels of such targets would be subject to changes in NNS availability. In this respect, NNS would behave like a class-specific transcription factor to fine-tune the expression of target genes.

## Acknowledgements

We would like to thank Domenico Libri, Mordechai (Motti) Choder, Ramon Grima and Julien Hurbain for critically reading the manuscript and providing valuable feedback. We would like to thank Ivan Clark, Alán Muñoz González, Lori Koch, Tessa Moses, Roopesh Krishnankutty, Martin Waterfall, David Kelly, Toni McHugh, and Sam Ranasinghe for their valuable discussions, helpful advice, and technical assistance. This work was supported by a Medical Research Council Non-Clinical Senior Research Fellowship (MR/R008205/1) to S.G., a Wellcome Trust PhD training fellowship (224084/Z/21/Z) and a Microbiology Society Research Visit Grant (GA003916) awarded to S.E-S., a Leverhulme Trust Grant (RPG-2018-004) and a Biotechnology and Biological Sciences Research Council Grant (BB/R001359/1) to P.S.S., and a Erasmus+ traineeship to T.W.

The metabolomics analyses were carried out by the EdinOmics research facility (RRID: SCR_021838) at the University of Edinburgh. Mass spectrometry was performed at the Mass Spectrometry Facility of the Institute of Genetics and Cancer (IGC) of the University of Edinburgh. Flow cytometry data were generated within the Flow Cytometry and Cell Sorting Facility in the Ashworth Building at the University of Edinburgh. The facility is supported by funding from the Wellcome Trust and the University of Edinburgh. The GFP reporters were synthesised with the assistance of the Edinburgh Genome Foundry, an engineering biology research facility specialising in the modular, automated assembly of DNA constructs at the University of Edinburgh. Training for confocal imaging was performed in the Centre Optical Instrumentation Laboratory (COIL), which is supported by a Core Grant (203149) to the Wellcome Centre for Cell Biology at the University of Edinburgh, and images were acquired at the Medical Sciences Confocal Imaging Facility at University College London. The Flp-In™ kit and the BY4742 strain were gifts from David Tollervey and Adele Marston, respectively.

## Author contributions

Conceptualization: S.G., P.S.S., M.D. and S.E-S. Methodology: S.G., P.S.S., M.D. and S.E-S. Investigation: S.E-S., T.W., M.G., I.F., P.S.S. and S.G. Visualization: S.E-S., P.S.S. and S.G. Writing - original draft: S.E-S. and S.G. Writing - review and editing: all authors. Supervision and funding acquisition: S.E-S., M.D., P.S.S. and S.G.

## Supplementary information

### Supplementary figure legends

**Figure S1. RNA-binding profiles of Nab3 and Nrd1 in low raffinose medium, related to Figure 1.**

(A) Schematic overview of the CRAC protocol and our differential peak calling strategy.

(B) Western blot and autoradiogram showing retrieved Nab3, Nrd1 and Sen1 ribonucleoprotein complexes in a *PIC2-GFP* parental strain grown in low raffinose medium. Signals were compared to those of a negative control (uncross-linked sample) and a positive control (Nab3-HTF grown in high glucose medium).

**Figure S2. Cell size aberrations and cell cycle delays co-segregate with mutations in the Nab3 binding sites of *PIC2*, related to Figure 3.**

(A) Weighted means and SDs of the cell size values obtained for three independent biological comparisons of the *GFP* carriers and WT *PIC2* spores resulting from *PIC2-GFP* and mut-Nab3-BS backcrosses with the non-fluorescent BY4742. All values were normalised to those of the *GFP* carriers stemming from the mating of both reference strains (i.e., BY4742 x *PIC2-GFP*).

(B) Schematic representation of Nab3 and Nrd1 RNA binding sites in *PIC2-GFP* and the mutations introduced in the indicated ‘mut’ strains.

(C) Fold changes of *PIC2-GFP* mRNA in all mutants compared to the parental reference in low raffinose medium. Bar plots display means and SDs from three independent experiments.

(D) Representative traces of the forward light scattering distribution of inspected cells for *PIC2-GFP* and derived mutants based on their forward light scatter (FSC-A) value, which is proportional to their cell size.

(E) Weighted means and SDs of the median FSC-A values for three independent biological repeats were calculated for each strain and normalised to that of the parental *PIC2-GFP* strain.

(F) Representative flow cytometry traces of cells with fluorescently dyed DNA of *PIC2-GFP* and the derived mutant strains at mid-log phase (OD_600_ ∼0.5).

(G) Bar plots displaying weighted averages and SDs of biological triplicates destined to quantify the population fraction of each strain within each cell cycle phase. *: p< 0.05,

**: p< 0.01 and ***: p< 0.001 and ‘ns’: p>0.05 (unpaired t-test).

**Figure S3. Altering expression levels of *PIC2* and *SLC25A3* in yeast and HEK 293 models, related to Figure 4.**

(A) Fold changes of *PIC2* mRNA in the *pTEF1-PIC2* and pic2Δ mutants compared to their parental strain. The bar plot displays averages, and error bars indicate SDs for three independent replicates.

(B) Log2-transformed fold changes of the population fraction of tested strains (i.e., WT, *pTEF1-PIC2*, and *pic2*Δ) against a fluorescent reference. Values shown are averages and SDs from biological triplicates and were normalised to the fold changes observed in the parental strain.

(C) Western blot comparing the levels of SLC25A3 after inducing its overexpression with increasing concentrations of doxycycline (Dox). The signal of SLC25A3 was normalised to that of GAPDH, a housekeeping protein with a higher molecular weight (kDa), in the same HEK 293 sample.

(D) Relative intensities of the SLC25A3 band with respect to that of GAPDH within the same lane. The bar plot shows averages, and error bars represent the SDs of two biological repeats. 0.2 µg/mL doxycycline was used to induce SLC25A3.

(E) Fold changes of *SLC25A3* mRNA abundance in HEK 293 cell cultures treated with 0.2 µg/mL doxycycline compared to uninduced cells. The bar plot shows averages and SDs for three biological repeats. *: p< 0.05, **: p< 0.01 and ***: p< 0.001 and ‘ns’: p>0.05 (unpaired t-test).

**Figure S4. Depleting Nab3 from the nucleus hampers cell cycle progression, related to Figure 5.**

(A) Representative flow cytometry traces of the forward light scattering distribution of inspected cells for *NAB3-FRB* and its parental WT strain based on their forward light scatter (FSC-A) value, which is proportional to their cell size.

(B) Representative flow cytometry traces of the DNA content of *NAB3-FRB* and its parental WT strain at several time points after rapamycin addition. Traces are representative of three independent biological repeats combined to generate the values shown in Figure 5C.

**Figure S5. Abolishing Nab3 binding to *PIC2* dampens Nrd1 binding to differentially expressed NNS targets such as *DUR3*, related to Figure 6.**

Visualisation of the raw reads obtained in Nab3 and Nrd1 CRAC and RNA-sequencing in the NNS co-regulated mRNA target *DUR3* in the *PIC2-GFP*, mut-Nab3-BS and mut-NNS-BS strains. To compensate for differences in library coverage, the raw reads shown in the y-axes were adjusted to display similar signals in neighbouring peaks which had not been deemed differentially bound by the DBPeaks package.

**Figure S6. Gene expression changes caused by transcriptome-wide redistribution of Nrd1 influence the cellular metabolic flux, related to Figure 6.**

(A) Dot plot summarising the average relative abundances and associated SDs of metabolites identified as differentially abundant in the Nab3 mutant (mut-Nab3-BS) across three independent biological comparisons with *PIC2-GFP*. Statistical analyses revealed that these differentially abundant metabolites were enriched in intermediates or products of the pyrimidine synthesis pathway, which is outlined in (B).

(B) Schematic overview of the pyrimidine synthesis pathway and Pic2 function in transporting phosphate and copper across the outer and inner mitochondrial membranes (OUM and IMM, respectively). The diagram underscores that, in *S. cerevisiae*, the pyrimidine synthesis biochemical route is not physiologically linked to the cellular function of Pic2, which is also depicted in the diagram. Whilst the metabolites and enzymes in pink font are upregulated in mut-Nab3-BS, their purple counterparts are significantly more abundant in the parental strain. IMS denotes intermembrane space.

## STAR Methods

### Key Resources Table

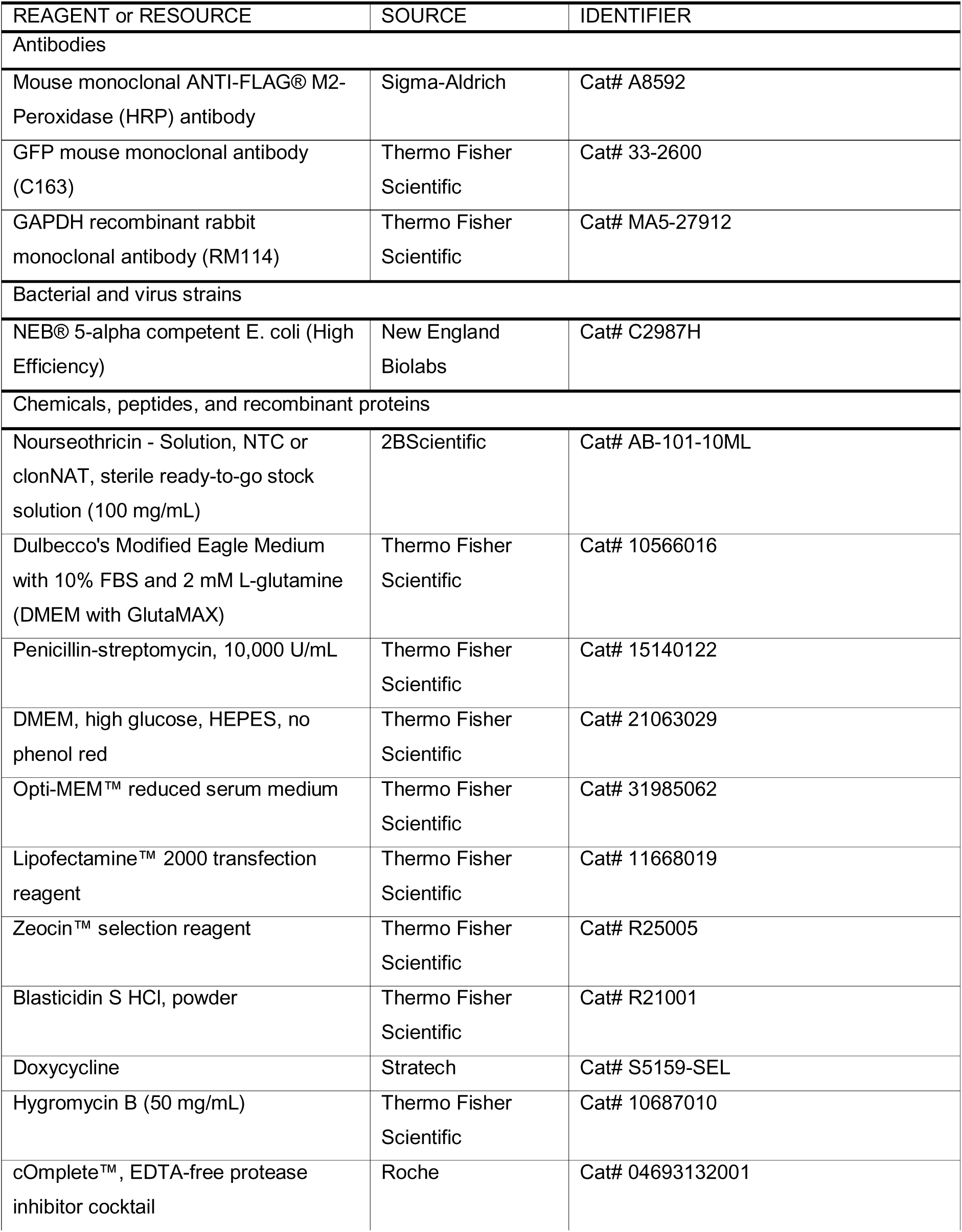
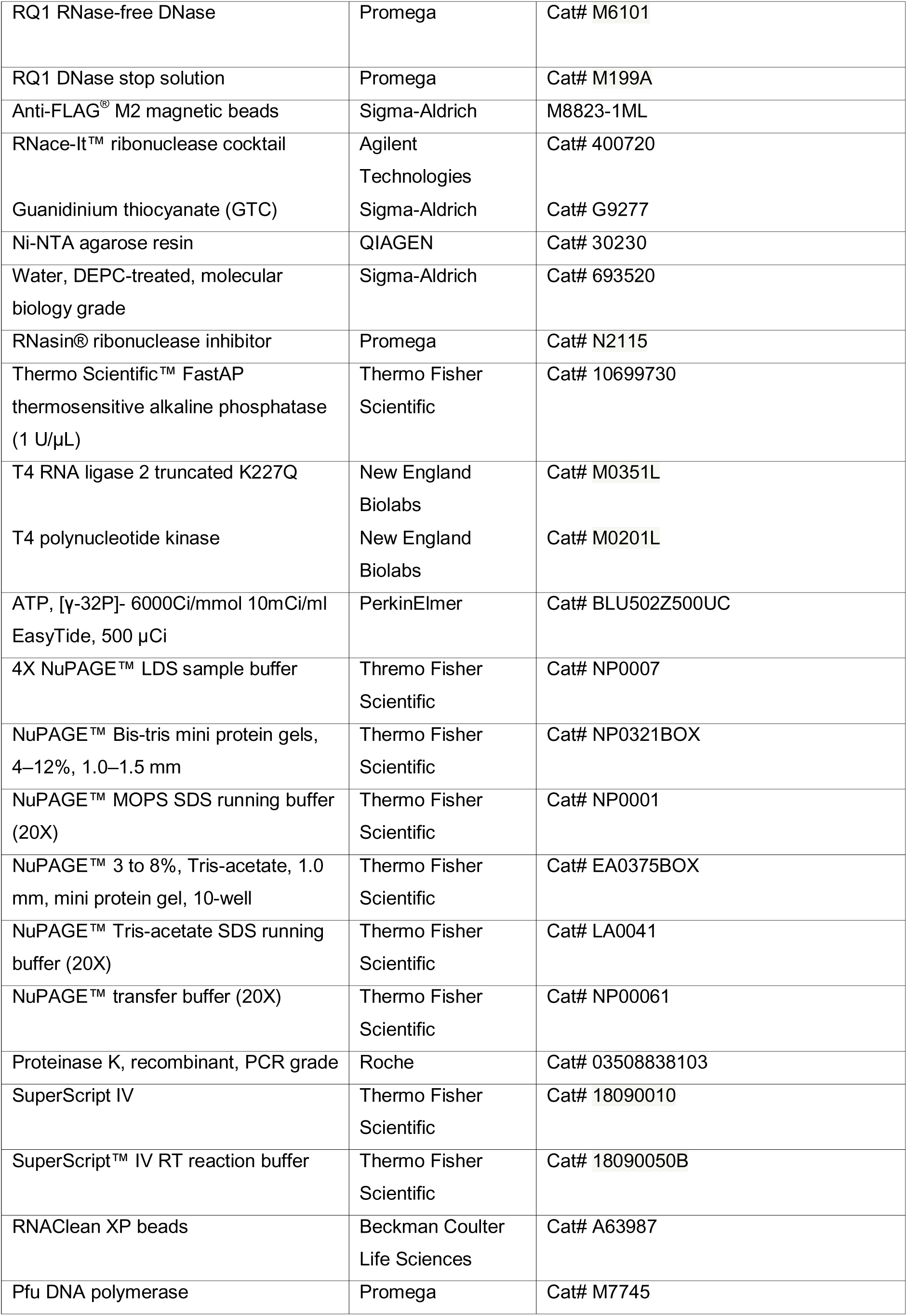
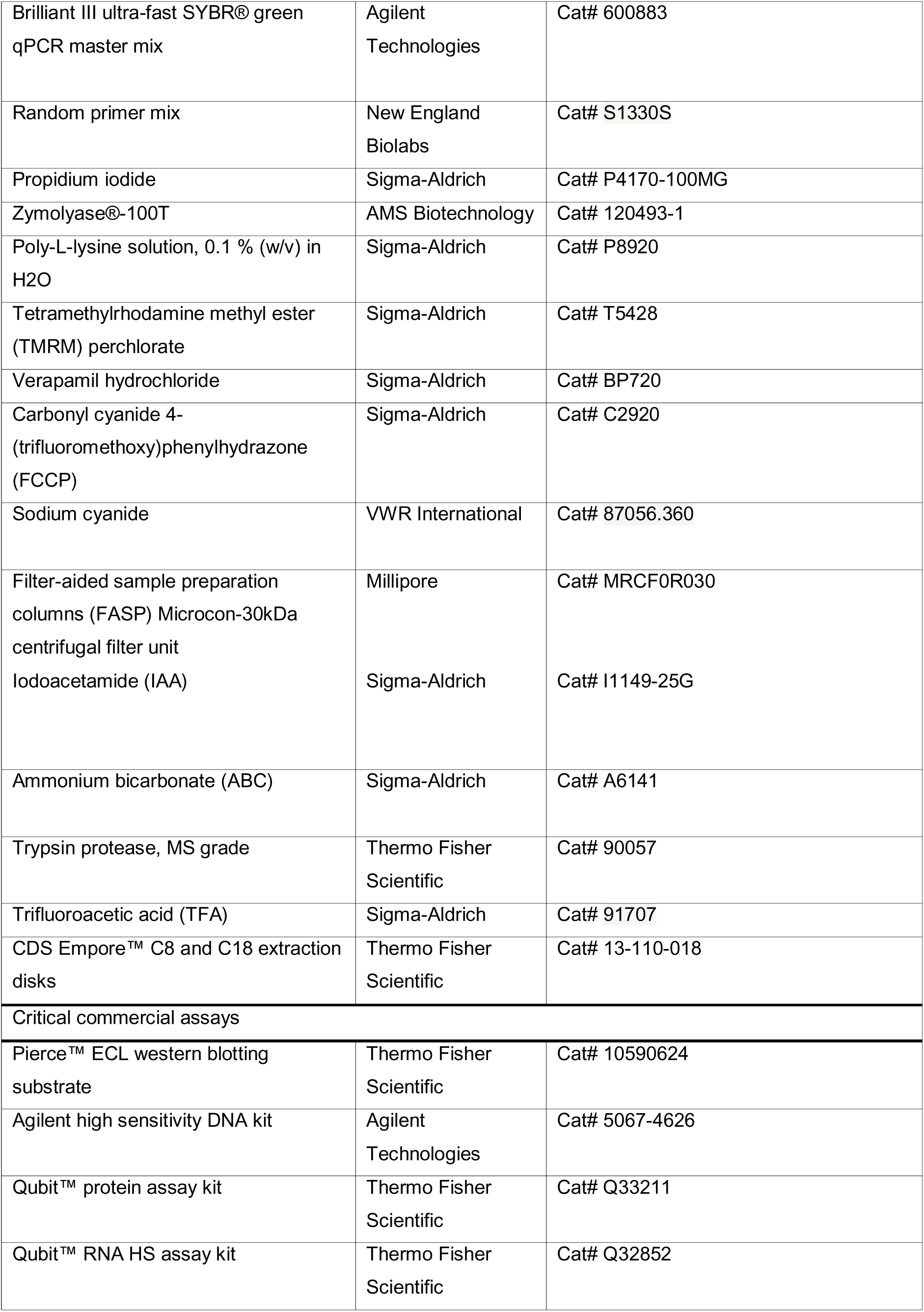
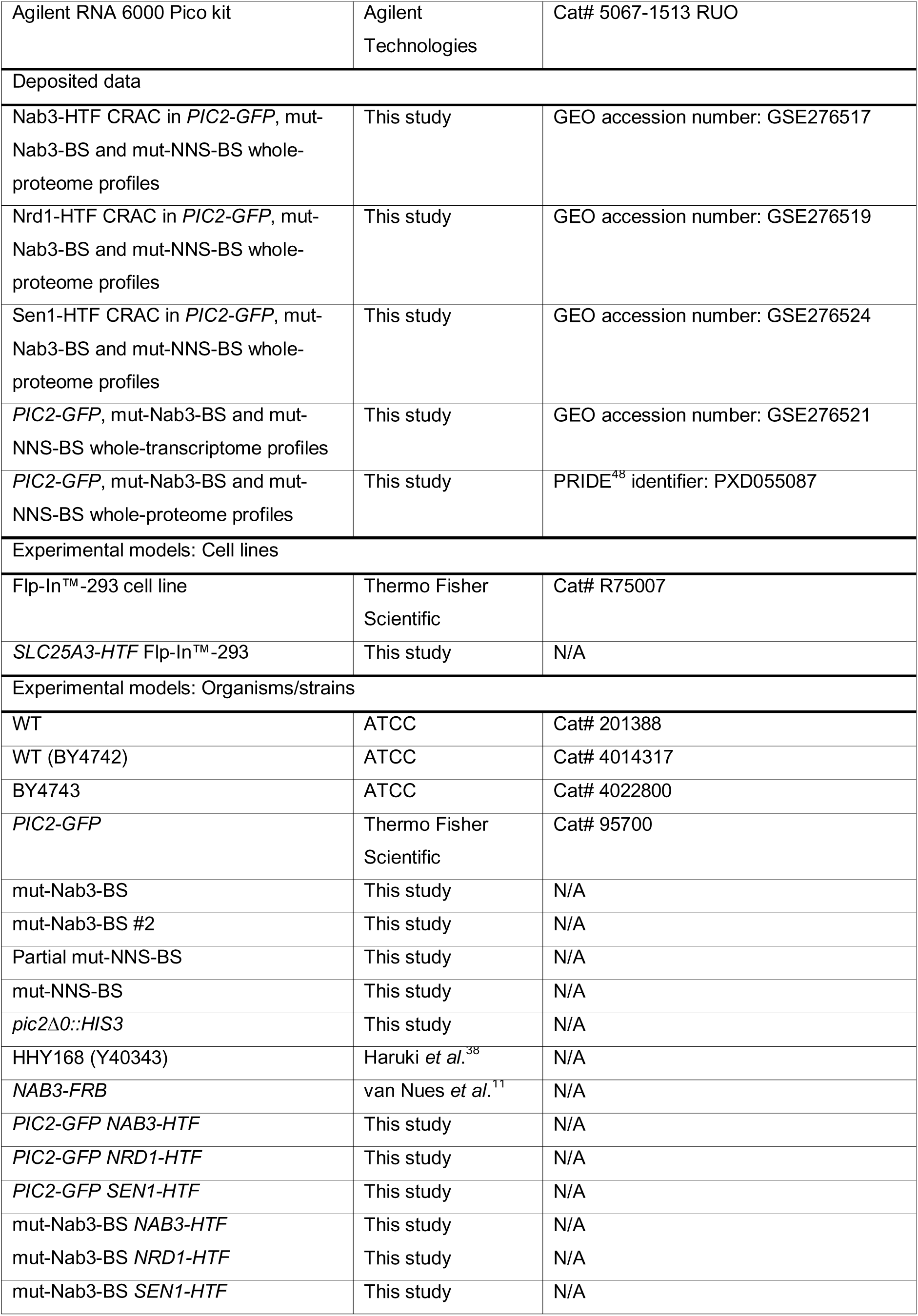
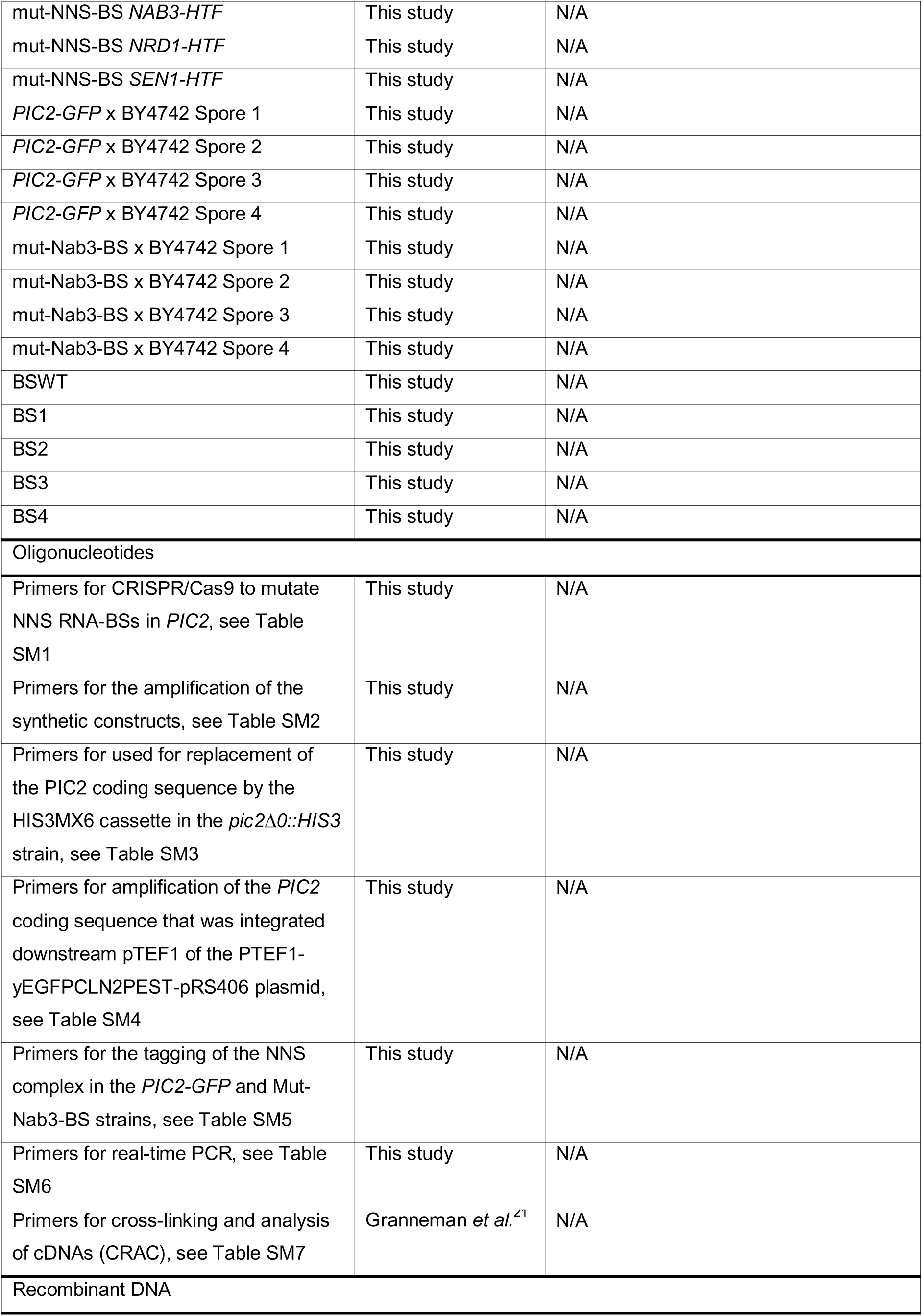
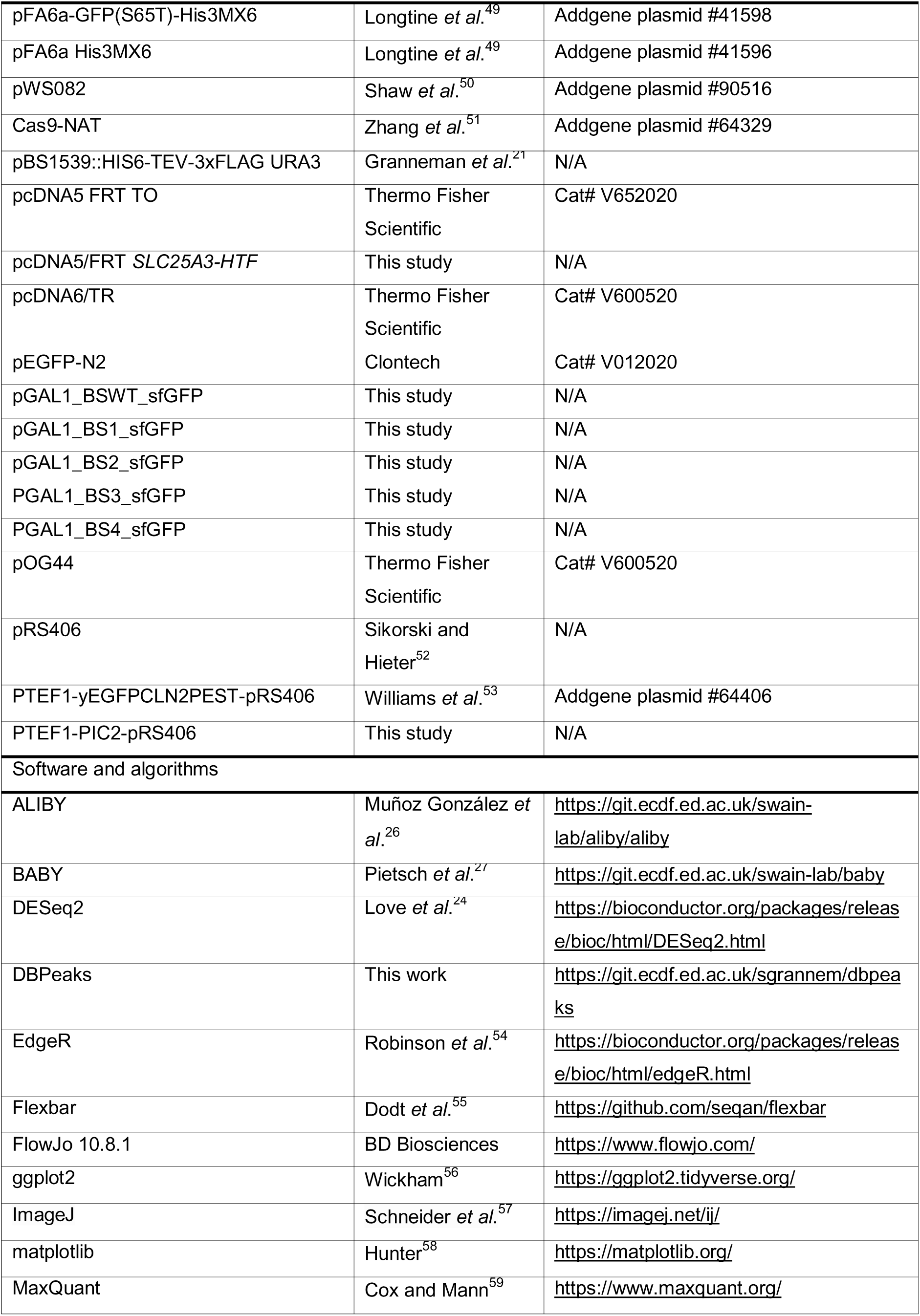
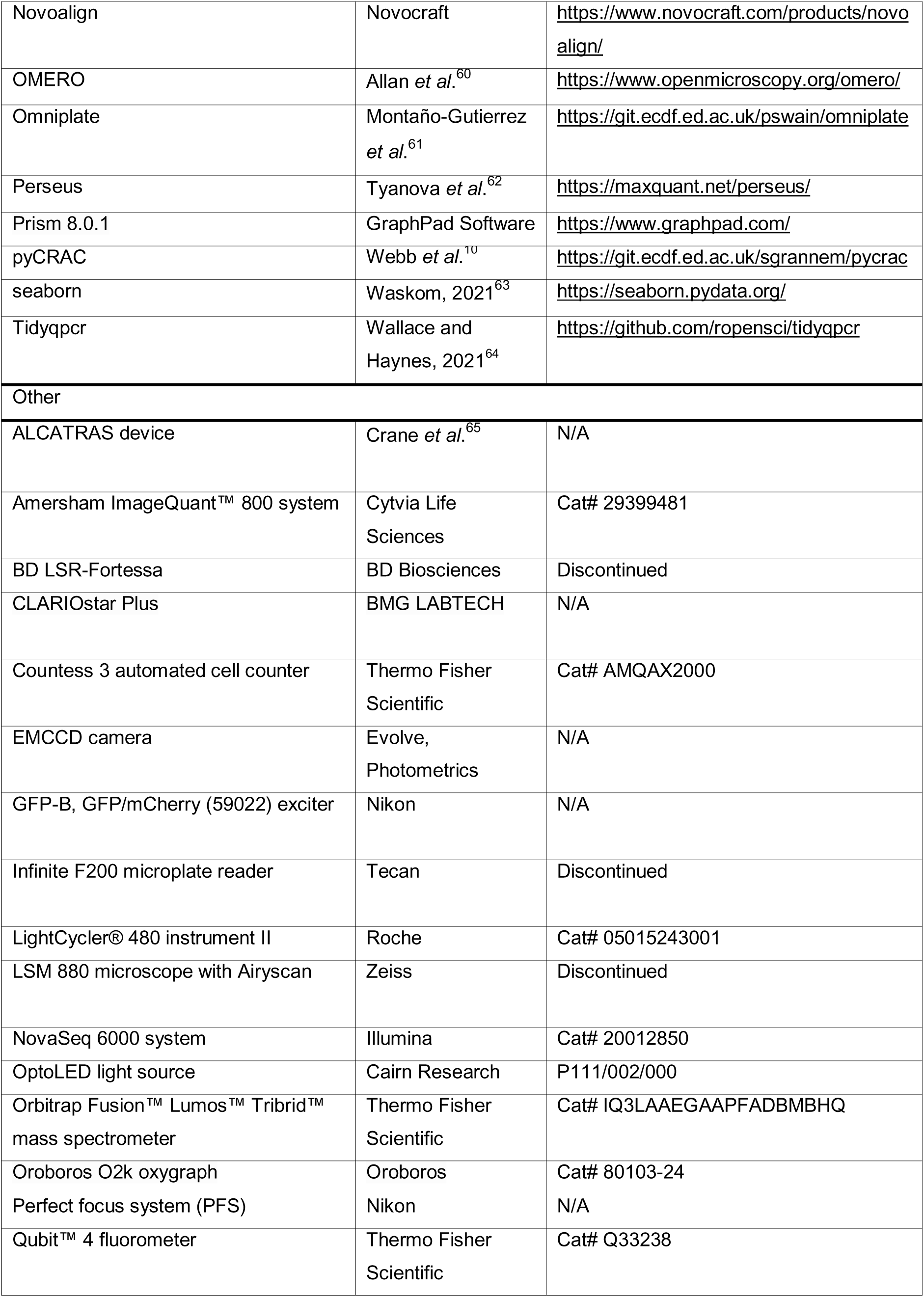
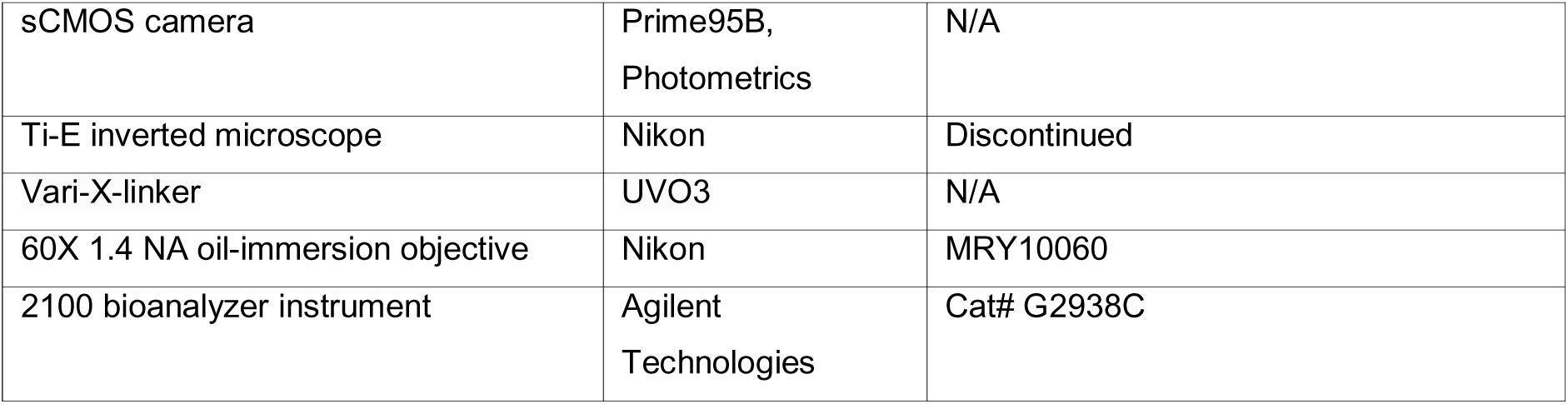

### Data availability

The raw mass spectrometry proteomic data and accompanying files have been deposited at the ProteomeXchange Consortium via the PRIDE^58^ partner repository. The accession number is PXD055087, also listed in the preceding Key Resources Table. The raw and processed RNA-sequencing have been deposited in NCBI Gene Expression Omnibus (GEO) under accession number GSE276521. Sequencing and peak-calling outputs for the CRAC libraries derived from Nab3, Nrd1 and Sen1 have also been uploaded to NCBI GEO under accession numbers GSE276517, GSE276519 and GSE276524, respectively.

### Contact for reagent and resource sharing

Further information and requests for resources and reagents should be directed to and will be fulfilled by the Lead Contact, Sander Granneman

### Method details

#### Yeast strain construction

All the strains engineered in this study are derived from the BY4741 reference^66^ and listed in the key resources table.

##### Homologous recombination

PCR-based modification was employed to fuse *GFP* downstream *PIC2*, to integrate HIS6-TEV-3xFLAG (HTF) epitopes distally to *NAB3*, *NRD1* and *SEN1*; and to delete *PIC2* in the *pic2*Δ*0::HIS3* mutant according to widely used protocols^49,67^. The coding sequence of *GFP*, the His6-TEV-3xFLAG (HTF) tag and the *HIS3* cassette were amplified from the pFA6a-GFP(S65T)-His3MX6^49^, pBS1539::HIS6-TEV-3xFLAG URA3^21^ and pFA6a His3MX6^49^ vectors respectively.

##### CRISPR/Cas9

A previously described CRISPR/Cas9 gene editing approach^68^ was applied to induce mutations in the promoter region of *PIC2.* The primers used are listed in Table SM1.

##### Modular cloning

Plasmids encoding the synthetic reporter constructs were designed based on the reported modular cloning (MoClo) system^69^ and adjusted to include the part containing the binding sites of Nab3 and Nrd1 as well as the corresponding spacer sequences. Parts were synthesised and assembled by the Edinburgh Genome Foundry.

##### Plasmid construction

The pTEF1-yEGFPCLN2PEST-pRS406^53^ backbone was cleaved by EcoRI/AscI to allow the integration of fragments containing the sequence of *PIC2,* which were amplified by primers containing MfeI and AscI restriction sites. The recombinant plasmid-encoded *URA3* as an auxotrophic marker and was transformed into the BY4741 strain to yield the *pTEF1-PIC2* mutant. To ensure medium uniformity across samples and experiments, all other strains used in this study were transformed with the unaltered pRS406 plasmid^52^, which also codes for the same selectable marker.

To generate the pcDNA5/FRT *SLC25A3-HTF* plasmid*, SLC25A3* was fused to an HTF tag, and NotI and XhoI restriction sites were included immediately upstream and downstream of the desired construct to enable digestion and subsequent integration of the *SLC25A3-HTF* coding sequence between the CMV promoter and the BGH polyadenylation signal encoded in the pcDNA5/FRT TO vector. The codon sequence of the *SLC25A3-HTF* construct was optimised for *Homo sapiens* by GeneArt (Thermo Fisher Scientific) and inserted into a standard (pMX) vector encoding a kanamycin-resistance selection marker. The designed vector was supplied as part of the GeneSyn service provided by GeneArt (Thermo Fisher Scientific). All plasmids used in this study are listed in the key resources table.

#### Mammalian cell line generation

Flp-In™ T-REx™ HEK 293 cells were co-transfected with three different vectors: one encoding SLC25A3 with its accompanying FRT recombination site, pcDNA5/FRT SLC25A3-HTF; another one coding for the flipase recombinase, pOG44; and a third one containing the tetracycline repressor and a blasticidin-resistance gene for selection, pcDNA6/TR.

##### Flp-In™ T-REx™ HEK 293 transfection

Having depleted antibiotics from the medium one hour before transfection, we dissolved 5 µg of each plasmid in OptiMEM medium (Thermo Fisher Scientific) to a final volume of 1250 µL. Similarly, for each 15-cm sample dish to be transfected, 15 µL of lipofectamine 2000 (Thermo Fisher Scientific) were mixed with 1235 µL of OptiMEM. Both mixtures were then combined, mixed by pipetting and incubated at room temperature for at least 20 minutes to allow the formation of DNA-lipofectamine complexes that can penetrate the mammalian plasma membrane of the cells in a single plate. Cells were then kept at 37°C for 1 hour before exposing them to a final concentration of 15 µg/mL blasticidin and 100 µg/mL hygromycin within their growth medium.

#### Yeast growth conditions

Yeast cells were grown in synthetic complete (SC) medium containing or lacking uracil (SC - Ura). Yeast strains encoding an auxotrophic marker were selected by growth in SC medium lacking uracil or L-histidine. Throughout microplate reader fluorescence assays, yeast cells were grown in SC medium lacking folic acid and riboflavin (LoFlo) to minimise background fluorescence. While a concentration of 2% (w/v) of a given sugar was considered rich medium, a 1 % (w/v) concentration posed a nutritional deprivation challenge to which cells in culture must adapt. Since fermentable carbon sources such as glucose are known to downregulate respiration, raffinose was used to formulate the media used in all experiments of this study except for those investigating the synthetic expression system. In the latter case, a mixture of 1% (w/v) sucrose and 1% (w/v) galactose was used to induce expression of the reporter gene from the *GAL1* promoter. Given that the expression of some hexose transporters can be influenced by previous exposure to fermentable sugars such as glucose^16,17^, all strains were pre-cultured in medium containing 2% (w/v) pyruvate for 48 hours prior to the start of the experiment^70^. This pre-growth stage ensured that the expression profiles of the different strains had a homogeneous starting point.

All liquid cultures were incubated at 30°C with shaking at 190 rpm. To facilitate growth in plates, medium for these was prepared using 2% (w/v) glucose as a carbon source and solidified upon the addition of agar to a final concentration of 1% (w/v). Cells grown in Petri dishes were incubated at 30°C in a static incubator for 48 hours and kept at 4°C afterwards. Yeast peptone dextrose adenine (YPDA) rich medium was used for growing cells before transformation and freezing. For the selection of transformants during the CRISPR/Cas9 genetic modification procedure, cells were plated in YPDA medium supplemented with 1% (w/v) agar and nourseothricin (100 μg/mL). The cells used in all experiments were harvested at OD_600_ 0.5 except those used in CRAC, which were gathered at OD_600_ 1. For microscopy and polarographic quantification of oxygen consumption, cells were grown in 5 mL cultures. Cytometry experiments, as well as RNA and protein extractions, were performed on cells collected from 25 mL cultures. Nab3 depletion in the *NAB3-FRB* strain was achieved by supplementing the pertinent 25 mL cultures with 1 μg/mL of rapamycin (Sigma-Aldrich). CRAC samples were obtained from 500 mL (for Nab3 and Nrd1 libraries) or 1 L (for Sen1) SC-Ura cultures containing 1% (w/v) raffinose. RNA and protein extractions, as well as CRAC experiments, were performed on cells that were either pelleted by centrifugation (3202 rcf, 5 minutes) or filtered after cross-linking, snap frozen by submersion in liquid nitrogen and stored at −80°C.

#### Flp-In™ HEK 293 cell growth conditions

Cells were grown in the Dulbecco’s Modified Eagle Medium (DMEM) containing 10% (v/v) foetal bovine serum (FBS), 2 mM L-glutamine, potassium penicillin (100 units/mL) and streptomycin sulphate (100 μg/mL). While the unmodified parental Flp-In™ HEK 293 cell line was grown in the presence of Zeocin™ (15 µg/mL), blasticidin (15 µg/mL) and hygromycin B (100 µg/mL) were supplemented to the growth medium of transfected cells. Transfected Flp-In™ HEK 293 cells were grown in 6-well plates to a confluence of approximately 80-90% before the start of all phenotypic characterisation experiments, and overexpression of the inserted *SLC25A3-HTF* was induced by adding 0.2 µg/mL of doxycycline to the growth medium 24 hours prior to the start of the experiment. To determine the minimum concentration of doxycycline required for significant overexpression of SLC25A3-HTF, its abundance was estimated by western blot at a range of doxycycline concentrations recommended by the cell line manufacturer.

#### Measuring growth and oxidative stress tolerance in yeast and mammalian cells

Yeast cells from a culture in the mid-logarithmic growth phase were resuspended in fresh medium to a starting OD_600_ of 0.1. 200 µL were transferred to three separate wells of a UV-sterilised 96-well black non-treated polystyrene microplate (Thermo Fisher Scientific) and its accompanying lid (Grenier). Triplicate controls comprising the same volume of sterile medium were also included for each examined condition. Absorbance and fluorescence were monitored in Infinite F200 (Tecan) microplate readers.

For Flp-In™ HEK 293 cells, 0.5 mM of hydrogen peroxide or an equivalent volume of sterile water were added to tested and control populations, respectively. Trypan blue (Thermo Fisher Scientific) was used to measure the viability of cell populations immediately after exposure to hydrogen peroxide or water and 18 hours after their addition. Counting was performed in a Countess 3 automated cell counter (Thermo Fisher Scientific). Two technical measurements were performed for each of the three biological assays that were completed per cell line and condition.

#### RNA extractions

Cells were thawed on ice and denatured by the addition of 200 μL of guanidinium thiocyanate (GTC) acid phenol mix. While mammalian cell lysis was achieved by vortexing at maximum speed twice for 30 seconds and incubation in a thermoblock for 5 minutes at 30° with shaking at 300 rpm, yeast cell lysis was performed by adding 400 μL of glass beads to each sample and vortexing all tubes at full speed for five 1-minute intervals separated by four 1-minute incubations on ice. An additional 1.5 mL of GTC phenol mix was then added to the latter samples. Yeast samples were incubated at 65°C for 10 minutes before being placed on ice for another 10 min. 800 μL and 100 µL of sodium acetate mix (3.3 mL 3 M NaOAc pH5.2, 0.2 mL 0.5M EDTA pH 8, 1mL 1M Tris-HCl pH 8, water to 100 mL) and 1.5 mL and 200 µL chloroform were pipetted into yeast and mammalian extracts respectively before vortexing each lysate at maximum speed for 5 seconds. Samples were centrifuged at 4°C (10621 rcf, 30 minutes), and the resulting aqueous phase was transferred into a new tube containing an equivalent volume of Phenol:Chloroform:Isoamyl alcohol mix. The mixtures were vortexed vigorously and centrifuged at 10621 rcf for 5 minutes. Once more, the aqueous phase of each sample was mixed with an equal volume of Chloroform:Isoamyl alcohol and vortexed at maximum speed before centrifugation at 10621 rcf for 5 minutes. Finally, the aqueous phase of each sample was relocated to tubes containing 3 volumes of 96% ethanol. The mix was vortexed vigorously and preserved at −80°C for at least 30 minutes or at −20°C overnight. Following ethanol precipitation, samples were spun at 10621 rcf for 30 minutes at 4°C. After discarding the resulting supernatants, 70% ethanol was pipetted into each tube. After centrifuging samples at 10621 rcf for 5 minutes at 4°C, ethanol was removed, and pellets were left to air-dry for approximately 5 minutes before resuspension in DEPC water. Finally, RNA concentrations were measured in the Qubit™ 4 fluorometer (Thermo Fisher Scientific) using the Qubit™ RNA HS assay kit (Thermo Fisher Scientific).

#### Real-time quantitative reverse transcription PCR

RNA extracts were purified by DNase treatment with RQ1 RNase-Free DNase (Promega) and reverse-transcribed using SuperScript™ IV reverse transcriptase and random primer mix (New England Biolabs). Samples were mixed with Brilliant III ultra-fast SYBR® green qPCR master mix (Agilent Technologies) and amplified within a 384-well plate (Roche) in LightCycler® 480 qPCR device (Roche). Three reference genes were inspected in every assay: while we used *ACT1*, *ALG9* and *PGK1* for yeast samples, we monitored *ACTB*, *GAPDH*, *RPL39* in mammalian ones. Sequences for all RT-qPCR primers are outlined in Table SM6.

#### Cross-linking and analysis of cDNAs (CRAC)

When SC-Ura cultures containing 1% (w/v) raffinose reached an OD_600_ of 1, cells were UV cross-linked (254□nm; 500□mJ/cm^2^) in the Vari-X-linker (UVO_3_;^11,20^) and rapidly filtered on 0.8 μm membranes (Merck) using a filtration device connected to a vacuum pump^11,20^. Membranes were collected in 50 mL tubes, flash-frozen by submersion into liquid nitrogen and stored at - 80°C until the day of the experiment.

On the day of the experiment, pellets were resuspended in a volume of lysis buffer (150□mM NaCl, 50□mM Tris pH 7.8, 0.1% (v/v) Nonidet P-40, 1 EDTA-free protease inhibitor cocktail (Roche) per 50 mL and 5□mM β-mercaptoethanol) corresponding to twice their mass. Lysates were then transferred to fresh 50 mL tubes, where they were mixed with 2 volumes of 0.5 mm zirconia beads (Thistle Scientific) and vortexed at maximum speed for five 1-minute intervals alternated with 1-minute incubations on ice. An additional volume of lysis buffer was pipetted into each sample before partitioning the cell debris by centrifugation at 4°C (3202 rcf, 15 minutes) and spinning down lighter impurities in an additional centrifugation step (10621 rcf, 20 minutes) at 4°C. Immunoprecipitation of the labelled protein from the supernatant was achieved during a two-hour incubation of the sample with 75□µL of anti-FLAG^®^ M2 magnetic beads (Sigma-Aldrich, M8823-5ML), which had been previously washed three times with 1□mL TN150 buffer. Immobilised beads underwent three 5-minute washes with TN1000 buffer (50□mM Tris pH 7.8, 1□M NaCl, 0.1% (v/v) Nonidet P-40 and 5□mM β-mercaptoethanol) and three additional 5-minute washes in 2□mL TN150 buffer (150□mM NaCl, 50□mM Tris pH 7.8, 0.1% (v/v) Nonidet P-40 and 5 mM β-mercaptoethanol). Afterwards, beads were resuspended in 550 µL of TN150 and incubated with 1 µL of a 1:100 dilution of RNace-It™ for exactly 5□minutes at 37□°C. To prevent overdigestion of co-immunoprecipitated transcripts and elute the bait proteins from the anti-FLAG beads, the solution was mixed with 0.4□g of guanidium hydrochloride (Gu-HCl) and the necessary volume of NaCl and imidazole (pH 8.0) to reach a final concentration of 300□mM and 10□mM respectively. After removing the beads, samples were pipetted onto 50□µL of Ni-NTA agarose resin (Qiagen) equilibrated with wash buffer I (6□M Gu-HCl, 10□mM imidazole, 300□mM NaCl, 50□mM Tris-HCl pH 7.8, 0.1% (v/v) Nonidet P-40 and 5□mM β-mercaptoethanol) and incubated at 4□°C overnight while rotating at 12 rpm.

Upon Ni-NTA binding, beads were transferred to Pierce columns (Thermo Fisher Scientific) and washed twice with 500□µL of wash buffer I before undergoing three additional washes of 500□µL with NP-PNK buffer (10□mM MgCl_2_, 50□mM Tris-HCl pH 7.8, 0.1% (v/v) Nonidet P-40, 5□mM β-mercaptoethanol). Alkaline phosphatase treatment was performed by resuspending immunoprecipitated RNA-protein conjugates in 80□µL of NP-PNK buffer containing 4 units of FastAP thermosensitive alkaline phosphatase (Thermo Fisher Scientific) and 80 units of RNasin® ribonuclease inhibitor (Promega) and incubating the samples at 37□°C for 1 hour in the case of Nab3 and Nrd1, or 15 minutes for Sen1. Dephosphorylation was stopped upon the addition of 500□µL of wash buffer. Having equilibrated the Ni-NTA agarose with three washes of 500□µL with NP-PNK, we performed adapter ligation by resuspending the beads 80□µL of reaction mix containing 0.6□µM of the applicable App-PE 3′ adapter (Table SM7), 30 units of T4 RNA ligase 2 truncated K227Q (New England Biolabs), 60 units RNasin® (Promega) and 10% (w/v) polyethylene glycol 8000 (PEG 8000). The reaction took place at 25□°C for six hours for Nab3 and Nrd1 CRAC and for three hours during Sen1 CRAC.

After washing once with 500□µL of wash buffer I and three times with 500□µL of NP-PNK buffer, we 5’end radiolabelled co-immunoprecipitated RNAs by incubating the Ni-NTA resin with 60□µL of NP-PNK buffer containing 30 µCi ^32^P-γATP (PerkinElmer) and 30 units of T4 polynucleotide kinase (New England Biolabs). After a 40-minute incubation of the reaction at 37°C, ATP (Roche) was supplemented to a final concentration of 1□mM, and the reaction continued for another 20 minutes. The reaction was stopped by washing three times with 500□µL of wash buffer I and three 500□µL washes of NP-PNK buffer followed before the addition of the 5′ linker ligation mix. The reaction took place in a volume of 80 µL of NP-PNK buffer supplemented with 1.25□µM of the fitting adapter, 40 units of T4 RNA ligase 1 (New England Biolabs), 80 units of RNasin® and 10□mM ATP (Roche). 5’ adapter ligation was left to proceed overnight at 16□°C for all bait proteins. The remaining steps of the protocol were executed as in previous reports^11,71^. After pooling suitable libraries together, the samples and their corresponding traces were submitted to Novogene, where paired-end sequencing of 150 bp-reads was performed in a NovaSeq 6000 system (Illumina).

#### Western blotting

Yeast and Flp-In™ HEK 293 cell lysates were prepared by addition of 2 volumes of lysis buffer (50 mM Tris-HCl pH 7.8, 150 mM NaCl, 0.1% (v/v) Nonidet P-40, 1 cOmplete™, EDTA-free Protease Inhibitor Cocktail (Roche) per 50 mL and 5□mM β-mercaptoethanol) to 1 g of cell pellet. Whereas no mechanical force was required for mammalian cell homogenisation, yeast samples were combined with 0.5 mm zirconia beads (Thistle Scientific) before being vortexed at full speed for five 1-minute intervals followed by four 1-minute incubations on ice. Cell debris was removed by centrifugation at 10621 rcf for 20 minutes. The protein concentration of each supernatant was measured using the Qubit™ 4 fluorometer (Thermo Fisher Scientific) and normalised to that of the most diluted sample. Protein analysis occurred in NuPAGE™ (Thermo Fisher Scientific) precast polyacrylamide gels. Whilst lysates assigned to Pic2-GFP, Nrd1-HTF, Nab3-HTF and SLC25A3-HTF detection were separated in 4% to 12% gradient Bis-Tris gels with MOPS running buffer (Thermo Fisher Scientific), samples designated to the inspection Sen1-HTF were partitioned in 3% to 8% gradient Tris-acetate gels with Tris-acetate running buffer (Thermo Fisher Scientific). All electrophoreses ran for 1.5 hours at 150 V. Proteins were transferred onto 0.2 μm nitrocellulose membrane and immunoblotting was performed using a 1:500 dilution of anti-GFP (for Pic2-GFP), a 1:1000 dilution of anti-GAPDH (for the loading control, GAPDH) and a 1:5000 dilution of a monoclonal anti-FLAG® M2-Peroxidase (HRP) antibody (Sigma-Aldrich, A8592; for HTF-fused proteins). Incubation with primary antibodies occurred overnight at 4°C. While the anti-FLAG antibody was directly imaged after chemiluminescence with the Pierce™ enhanced chemiluminescence (ECL) kit (Thermo Fisher Scientific), the anti-GFP and anti-GAPDH antibodies underwent a 2-hour hybridisation to 1:5000 dilutions of anti-mouse IRDye 800CW (LI-COR Biosciences) and goat anti-rabbit IRDye® 680RD (LI-COR Biosciences) secondary ones at room temperature. HTF-tagged proteins were visualised by applying the automatic chemiluminescence exposure settings in the Amersham ImageQuant™ 800 system (Cytvia Life Sciences). Membranes requiring fluorescence imaging were exposed to the IR long (775 nm) and IR short (660 nm) channels of the same device for 15 seconds and 5 minutes, respectively.

#### Flow cytometry

##### DNA quantification for cell cycle analysis

5 mL of cells from mid-exponential-phase cultures were pelleted and immediately fixed by the addition of an equal volume of ice-cold 70% ethanol and an overnight incubation at −20°C. On the next day, cells were permeabilised with 800 µL of 50 mM sodium citrate buffer (pH 7.2) and a 10-minute incubation at room temperature. This step was repeated once more before adding RNace-It™ ribonuclease cocktail (i.e., a mixture of RNase A and RNase T1; Agilent Technologies) to a final concentration of 0.1 mg/mL (i.e. 15 µL of 2 mg/mL stock) and incubating the samples overnight at 37°C. Following RNA degradation, proteins were digested upon the addition of 10 µL of 20 mg/mL proteinase K (Roche) and subsequent incubation at 55°C for 2 hours. Afterwards, cell aggregates were disrupted by sonication (5 seconds, 25 µm). At this stage, samples could be stored for up to 1 month at 4°C or stained with 50 μg/mL propidium iodide (Sigma-Aldrich) for flow cytometry. In the latter case, 200 µL of each resulting sample were pipetted into individual wells of a clear 96-well plate (Thermo Fisher Scientific) for inspection in a BD LSR-Fortessa (BD Biosciences) at the Flow Cytometry Facility for the School of Biological Sciences of the University of Edinburgh. 10,000 events were recorded per acquisition.

##### Fitness assay

A volume holding an equivalent to OD_600_ 0.0005 of the tested strain was transferred into 5 mL of fresh medium before adding the red fluorescent strain (Ura7-mCherry) at a final identical OD_600_. Cells were then left to compete in the medium overnight. After 18 hours, 1 mL of culture was pelleted (3202 rcf, 5 minutes) and resuspended in PBS prior to inspection in a BD LSR-Fortessa (BD Biosciences). A minimum of 20,000 events were recorded per competition culture. A volume corresponding to OD_600_ 0.0005 of the mid-logarithmic-phase culture was transferred to 5 mL of the same sterile medium to continue monitoring the competition throughout time. Subsequent samples were gathered at 36- and 60-hours post-inoculation for analogous flow cytometry acquisitions.

#### Mating assay

*PIC2-GFP*, mut-Nab3-BS and mut-NNS-BS (MATa) were mixed with a BY4742 (MATtα) in YPDA plates and incubated at 30°C. The following day, single colonies were re-streaked into fresh YPDA plates and stored at 30°C to allow growth overnight. Mating tester plates were then prepared to screen for diploids, which were able to grow independently from the mating factor (a or α) which had been used to coat such plates. Potential diploids underwent sporulation upon re-streaking in SPO sporulation medium (0.3% (w/v) potassium acetate, pH 7). Throughout an incubation at 30°C for up to 3 days, plates were checked regularly for tetrads, and once they were identified, these were collected for digestion. To this end, cell samples were resuspended in 20 μL of digestion solution (1 mg/mL of zymolase (AMS Biotechnology) dissolved in 1 M sorbitol) and subsequently incubated at room temperature for 8 minutes. To stop the reaction, 1 mL of sterile water was added to each sample. 20 μL of the resulting solution were pipetted on the border of a fresh YPDA plate and left to slide through its diameter before being air-dried. Once all the volume of the digestion solution had evaporated, a minimum of 8 tetrads were dissected onto each half of the plate. Dissection plates were then incubated at 30°C overnight before proceeding with a mating-type check of the sister spores within each tetrad by colony PCR or microscopy inspection. Dissections yielding an a to α ratio of 2:2 were deemed successful, and hence, we proceeded to quantify the cell size of their constituent spores.

#### Quantification of mitochondrial membrane potential with confocal microscopy

Mitochondrial membrane potential was assessed using tetramethyl rhodamine, methyl ester (TMRM; Sigma-Aldrich), a cell-permeable, positively charged, red fluorescent dye that accumulates in mitochondria proportionally to their activity. TMRM was added to mid-exponential cultures at a final concentration of 50 nM. After a 30°C with shaking at 190 rpm, 100 µL of cell solution were then pipetted into the wells of a black 96-well plate with bottom glass (Grenier), which had been pre-coated with 0.1% (w/v) poly-L-lysine (Sigma-Aldrich). Poly-L-lysine effectively allowed temporary immobilisation of some cells during live imaging.

Adherent Flp-In™ HEK 293 cells were grown directly in a 96-well plate with a bottom glass. Over 12,000 mammalian cells were seeded 4 days before the imaging was performed. Fresh DMEM and the relevant antibiotics were provided every 48 hours. Two days after seeding, fresh medium was replenished, and *SLC25A3* overexpression was induced with 0.2 µg/mL doxycycline 24 hours before the start of the experiment. On the day of the experiment, DMEM was removed, and cells were washed twice with 100 μL of PBS and finally incubated at 37°C in recording medium (phenol red-free DMEM (Thermo Fisher Scientific) supplemented with 10 mM glucose, 1 mM glutamine and 10 mM HEPES, pH 7.4). 30 minutes before imaging a given well, 25 nM of TMRM was added to the recording medium. To prevent membrane transporters from extruding the dye, cells were also treated with 25 μM of verapamil (Sigma-Aldrich), an efflux pump inhibitor.

Imaging was then performed in an LSM 880 microscope (Zeiss) at the Medical Sciences Building Confocal Imaging Facility at University College London. While yeast and mammalian samples were kept at 30°C or 37°C respectively inside an incubation chamber (Zeiss). For yeast acquisitions, TMRM was excited with a 561 nm argon laser with an output power of 0.2 mW, the pinhole was set at 2.67 arbitrary units (a.u.), and the detector gain was fixed at 850. Whereas red fluorescence parameters remained unaltered for all acquisitions, the detector gain for the T-PMT channel had to be adjusted to successfully view the outline of the cells in each sample. In both channels, 10 z-stacks were taken for TMRM images, with a spacing of 0.6 μM between slices. For Flp-In™ HEK 293 imaging, TMRM settings were slightly modified compared to those of yeast cells: the argon laser power was decreased to 0.1 mW to minimise radiation-induced stress in the cells, the pinhole was reduced to 1.62 a.u. to increase the resolution of the stained areas and, owing to the larger volume of human cells, the z-stacking parameters were changed to acquire 14 regularly spaced slices spanning 13 μm. To verify that the signal captured in the red fluorescence channel emerged solely from TMRM, we acquired control images of unstained yeast cells. Additionally, to confirm that TMRM was exclusively being sequestered by active mitochondria, we re-imaged stained samples in which mitochondrial membrane potential was collapsed upon the addition of carbonyl cyanide-p-trifluoromethoxy phenylhydrazone (FCCP; Sigma-Aldrich), a powerful uncoupler of oxidative phosphorylation.

#### Polarographic quantification of oxygen consumption rates

Oxygen consumption rates were measured using a high-resolution O2K respirometer (Oroboros). Cells from mid-exponential cultures were resuspended into fresh sterile medium to reach a final OD_600_ of 0.1. 2 mL of these cell solutions were injected into a pre-warmed chamber before recording basal respiration measurements at 30°C or 37°C for yeast or HEK293, respectively.

#### NADH fluorescence measurements

Fluorescence measurements were recorded using the OD_600_ absorbance and NADH fluorescence modes of the CLARIOstar Plus (BMG LABTECH) microplate reader. OD_600_ measurements were obtained to enable the normalisation of the NADH fluorescence values to cell numbers. Given that the nicotinamide moiety of NADPH would also fluorescence in this mode, the raw recording is, in principle, also dependent on the status of the NADP(H) pool. However, the concentration of NAD(H) is around 100 times larger than that of NADP(H)^72^, and the NADP pool is stably maintained in its reduced state (i.e., at least 20:1 NADPH to NADP)^73^. Therefore, even the raw measured fluorescence signal is broadly reflective of the state of the NAD(H) pool.

To compensate for the effect that emissions proceeding from NADPH may have in the NADH measurement, the resting fluorescence value was normalised to the fluorescence recordings in samples in which the NADH/NAD^+^ ratio reached its minimum and maximum, respectively^74^. While the basal measurement was taken in untreated cells, another set of cells was exposed to FCCP, which elicits maximal respiration and, consequently, drives full oxidation of the NAD pool to its NAD^+^ state. Thus, any NADH signal recorded upon the addition of FCCP was assumed to correspond to a background measurement for the NADH autofluorescence acquisitions. Conversely, another set of samples was treated with sodium cyanide (NaCN; VWR International), which blocks complex IV of the electron transport chain and impedes respiration. Even though NaCN-mediated inhibition of complex IV is not substrate-dependent, the physiological response of the cell to a shutdown of the electron transport chain is to increase the production of substrates that can be fed to the ETC to resume respiration. Consequently, the cell fully reduces its NAD pool, thereby yielding its maximum NADH fluorescence. NADH fluorescence values were recorded 2 minutes after the addition of the drugs. Due to their different susceptibility to FCCP and NaCN, yeast cells were exposed to 15 µM and 1.25 mM of these drugs, respectively, whereas Flp-In™ HEK 293 cells were treated with 1 µM of FCCP and 1 mM NaCN. Whereas yeast acquisitions were performed at 30°C, HEK 293 ones occurred at 37°C.

#### RNA sequencing

RNA extracts were treated with RQ1 RNase-Free DNase as per the manufacturer’s guidelines. The quality of the RNA samples was then inspected in an Agilent 2100 bioanalyzer (Agilent Technologies) using an RNA 6000 pico assay (Agilent Technologies) according to the manufacturer’s instructions. Samples with an RNA integrity number (RIN) score greater than 7 were submitted to Novogene for paired-end sequencing of 150 bp-reads in a NovaSeq 6000 system (Illumina). Before sequencing, samples underwent a rRNA depletion step, and libraries were generated using the TruSeq preparation protocol (Illumina).

#### Sample preparation for proteome profiling

Cell pellets were thawed and resuspended in 20 μL of 4% (w/v) SDS dissolved in 100 mM Tris-HCl pH 8 and incubated at 95°C for 30 minutes. Samples underwent three sonication cycles (10 seconds ‘on’, 5 seconds ‘off’) at an amplitude of 5 μm. Tris-HCl pH 8 was added to dilute SDS concentration to 1% (w/v). Additionally, dithiothreitol (DTT) was supplemented to a final concentration of 10 mM before incubating the lysates at 50°C for 30 minutes. Samples were then centrifuged at 20817 rcf for 15 minutes before measuring the protein concentration in the resulting supernatant. A volume holding 100 μg was aliquoted for each sample before adding urea to a final concentration of 6 M. Following mixing, denatured proteins were cleaned with filter-aided sample preparation columns (FASP, Microcon-30kDa centrifugal filter unit, Millipore, MRCF0R030). After centrifugation (14000 g, 40 minutes), the flow-through fraction was discarded, and the filter membrane was further washed with 200 μL of 8 M urea in 100 mM Tris-HCl pH 8. The proteins in the filter were then alkylated with 100 μL of 5 mM iodoacetamide (IAA, Sigma-Aldrich) dissolved in 8 M urea. Alkylation occurred at room temperature in the dark for 20 minutes. The filter membrane was then washed once with 100 μL of 8 M urea Tris-HCl pH 8 and twice with 100 μL of 50 mM ammonium bicarbonate (ABC, Sigma-Aldrich). Upon centrifugation (14000 g, 10 minutes) and subsequent removal of the flow-through fraction, 1□µg of mass spectrometry grade trypsin protease (Thermo Fisher Scientific) was diluted into 39□µL ABC and applied to the column of each membrane. All samples were incubated at 37□°C overnight until an additional 40 μL of ABC were added to each sample, and its peptides were collected by centrifugation (14000 g, 10 minutes). The peptide concentration was measured using a Qubit™ 4 fluorometer (Thermo Fisher Scientific) before samples were acidified to pH≤3 with the pertinent volume of 10% (v/v) trifluoroacetic acid (TFA) and desalted in C18 extraction filters (Thermo Fisher Scientific) contained within 200 µL microtips. Membranes were activated upon passing (14000 g, 10 minutes) of 15□µL methanol and equilibrated with a 50□µL-wash of 0.1% v/v TFA membrane. After drying their filters by centrifugation (14000 g, 10 minutes), tips were stored at −20°C immediately before mass spectrometry. Peptides were eluted with 40□µL 80% acetonitrile (ACN) in 0.1% v/v TFA and tandem mass spectrometry (MS/MS) analysis was performed on an Ultimate Ultra3000 chromatography system coupled with a Orbitrap Fusion™ Lumos™ Tribrid™ mass spectrometer (Thermo Fisher Scientific) at the Mass Spectrometry Facility of the Institute of Genetics and Cancer (IGC) of the University of Edinburgh using previously specified loading parameters^75^.

#### Whole metabolome profiling

500 μL of cultures in the mid-exponential growth stage were quenched by submersion into a dry ice/ethanol bath for 10 seconds. Samples were then spun down at 4°C (1000 g, 10 minutes) before discarding 150-450 μL from the top fraction. The remaining volume underwent an additional centrifugation (2500 g, 5 minutes) at 4°C. The resulting cell pellet was resuspended in a 10-fold volume of ice-cold HPLC-grade chloroform:methanol:water (1:3:1) mixture. An equivalent volume of 0.5 mm zirconia beads was added to the samples before they were vortexed at maximum speed for five 1-minute intervals spaced by four 1-minute incubations on ice. Lysates were then incubated at 4°C for 1 hour with shaking at 1200 rpm. Samples were subsequently vortexed at maximum speed for 5 minutes on a cooled mixer before being spun down (13000 g, 3 minutes) at 4°C to separate the cell debris from the metabolite-containing supernatants, which were stored at −80°C until they were analysed. Metabolite characterisation was performed by the EdinOmics facility at the University of Edinburgh following a previously reported methodology^76^. Statistical analysis was performed using the MetaboAnalyst 5.0^77^ web-based platform, and dot plots were generated in GraphPad Prism 8.1.

#### Quantitative analyses

##### Differential peak calling using DBPeaks

Raw sequencing outputs were returned as FASTQ files. Hence, after conversion into FASTA format, files were processed using the paired-end (PE) version of the pyCRAC pipeline^10,11^. This package demultiplexes barcoded reads and removes adapter sequences by applying the previously developed Flexbar tool^55^. Having been filtered based on their length and nucleotide quality values (i.e., PHRED scores), the remaining reads were gathered into BAM outputs and fed into the newly developed DBPeaks.py peak calling script (https://git.ecdf.ed.ac.uk/sgrannem/dbpeaks). Firstly, The `countReadsBam` function of the DBPeaks script employs the pyReadCounters tool from the pyCRAC package to generate GTF files with read counts and gene mapping locations. Next, the `getPeaks` function uses the `pyCalculateFDRs` script from the pyCRAC package to retrieve peaks enriched in each library relative to randomly distributed reads over the same genes. The resulting peak GTF files are then filtered by the `filterPeaks` function to remove peaks comprising fewer reads than the mean coverage of all the intervals in the dataset. Although we used mean, median or mean with one standard deviation can be selected as thresholds. Peak widths were normalised to a minimum length of 10 nucleotides with the `adjustPeakWidths` function. Afterwards, the `mergeGTFintervals` applied BEDTools^22^ to concatenate intervals or peaks that are present in all replicates. Subsequently, the `numberPeaks` function assigns a number to each filtered peak to prevent misidentification in downstream steps. The software then performs a pyReadCounters analysis on the BAM files using the new GTF file encompassing the numbered filtered peaks. We then gathered the hit tables outputted by pyReadCounters and combined them using the `mergeHittables` function. Finally, the `runDESeq` function automatically performs the statistical analysis to identify differential peaks using the DESeq2 pipeline^24^. By default, DESeq2^24^ returns results in a TXT file, so we developed two additional functions to extract the calculated fold change and incorporate it into the original peak file. Finally, the `getSignificantPeaks` function generates a new GTF file containing solely the peaks that were called significant by the DESeq2^24^.

##### Real-time quantitative PCR analyses

Raw Cq values were analysed using Tidyqpcr package^64^. The resulting fold change values in *PIC2* mRNA abundance in each mutant were individually compared to their corresponding parental strain using unpaired t-tests. Statistical analyses and plotting were performed in GraphPad Prism 8.1.

##### Microfluidics inverted microscopy image analyses

Experimental outputs were uploaded to an OMERO server^60^. To analyse the microscopy images, we used the ALIBY pipeline^26^, which automates cell segmenting, tracking and post-processing of microscopy time-lapse images. This package fetched the images from Omero and tracked the tiles constituting the traps present across time-lapse images of every position that was established during the experimental set-up. In doing so, the software accounted for any spatial drifting which may have occurred while the images were captured. The pipeline then uses the Birth Annotator for Budding Yeast (BABY) algorithm^27^ to create masks encircling the outline of the cells in the previously defined traps and subsequently tracking them across time points to generate cell lineages. Cellular outlines are used to delimit the acquisitions in the brightfield and GFP channels and, consequently, extract the area, budding events, and fluorescence intensity of each cell and to estimate the volume of each cell^27^. Having computed the average fluorescence in regions laying outside cell masks, we subtracted this background from the intensity values assigned to each cell in the same image. We corrected cells too for autofluorescence by subtracting the fluorescence of cells from an untagged wild-type strain. These untagged cells were matched to the fluorescent cells both by their volume and their time of measurement. We estimated the autofluorescence using the fluorescent cell’s volume and a LOWESS interpolation of the fluorescence versus volume relationship for untagged cells, recalculated in 15-minute intervals. Data was visualised using seaborn^63^ and matplotlib^58^ Python libraries.

##### Yeast growth and fluorescence analyses

Microplate reader data was analysed using the Omniplate software^61^, which processes time-series data raw outputs from microplate readers to quantify microbial growth. Omniplate blanked OD measurements by discounting absorbance signal from the wells containing media only and corrected for the non-linear relationship between OD_600_ measurements and cell count at higher absorbance values. The software then makes assumptions about the family of latent functions that can describe a population using a Gaussian-process-based algorithm^61^. In this way, Omniplate can predict errors in the estimated growth rates by fitting additional artificial replicates of the experimental data using bootstrap or iterative random re-sampling^78^. Having extracted the OD values were extracted from the triplicates for each sample, we plotted the average growth rate with an associated error corresponding to half of the interquartile range.

The `getstats` function computed the maximal growth rate that had been estimated based on the three technical replicates that were included for each sample^61^. To determine whether a strain was resistant or hypersensitive to oxidative challenges such as hydrogen peroxide, we calculated the logarithmic ratio of the maximal growth rate in medium containing 10 mM hydrogen peroxide to that of the medium lacking the reactive oxygen species. The resulting logarithmic fold changes could then be statistically compared using unpaired t-tests performed by GraphPad Prism 8.0.1, which was also used to generate the relevant bar plots.

For fluorescence assays, the software subtracted background fluorescent signals inherent to the media from the fluorescence readings of wells containing cells^61^. The resulting values were then corrected for autofluorescence, which was defined as the fluorescence signal emerging from an unmodified BY4741 parental strain. Final corrected fluorescence values were normalised to corrected OD_600_ measurements and plotted against time using Python’s seaborn package^63^.

##### Flow cytometry analyses

Experimental files were exported in their native FCS format and analysed in FlowJo 10.8.1. For cell size experiments, the normalised distribution of ungated FSC-A values was plotted in histograms. The weighted average and standard deviation were calculated for the three independent biological repeats that were assessed per cell type and condition. For cell cycle analyses and competition experiments, we introduced a gate to select for single-cell events and subsequently generated histograms of PE-Texas Red-A. To analyse DNA content in the cell cycle experiments, red fluorescence histograms were standardised to the mode of the events within all the repeats of the appropriate control strain and subsequently used to draw gates delimiting each cell cycle stage. The cell counts at each phase of the cell cycle were obtained in three independent biological repeats and used for statistical comparisons between strains. In the case of competition experiments, a gate was applied to delineate samples containing either non-red parental strain (i.e. BY4741) or the red reference (Ura7-mCherry) cells only. The counts of non-red and red cells within a competition culture were used to calculate the ratios of non-fluorescent to fluorescent cells. Given that each competition was performed in three independent repeats, a mean and standard deviation for that ratio could be determined for each strain. Non-fluorescent: fluorescent ratios and their errors were normalised to that of the pertinent reference strain and subsequently underwent a logarithmic transformation. Mutants were compared to their parental strain using unpaired t-tests ran in GraphPad Prism 8.1, which was also used to generate the summary histograms.

##### Confocal microscopy image analyses

Images were saved as CZI files and viewed with standard Image J^57^ in a default stack order. Channels were split, and regions of interest (ROIs) were drawn and added to the PTM channel. Having selected the cells that were examined, we saved ROIs as ROI files for re-evaluation. In the TMRM channel, the image’s stacks were z-projected to obtain the maximal intensity. The image’s threshold was then adjusted using the Otsu’s method for image segmentation, which is an algorithm for image analysis that converts fluorescence to grayscale, assigns an intensity value for every pixel and, thenceforth, partitions the latter into foreground and background based on a threshold value. All the parental replicates were assessed individually for their Otsu’s optimal threshold and the highest threshold value was then imposed as a common one to discriminate between stained mitochondria and background TMRM fluorescence in all remaining images of the derived strains.

To correct for the varying background fluorescence of individual images, we specified a ROI in a space without cells and subsequently measured its fluorescence on the TMRM channel without introducing a minimal threshold. This background fluorescence was then subtracted from the mean fluorescence for the maximum projection of each selected ROI in the same image. Corrected fluorescence was then normalised to the mean corrected fluorescence of the corresponding replicate of the parental strain for direct comparison. At least 50 cells from three independent biological repeats were examined for each strain and cell line. Unpaired t-tests and visualisation were performed in GraphPad Prism 8.1.

##### Respirometry analyses

Experimental traces were saved and exported in the native DLD format for analysis in the Oroboros Datlab 7 software (Oroboros). Having identified and demarcated sections on which the oxygen consumption signal remained relatively constant, we extracted the average value of these marks and used three biological replicates to calculate the mean and standard deviation for the basal respiration rates. Statistical significance was evaluated by unpaired t-tests performed in GraphPad 8.1, which was also used to assemble the bar plots showing the mean and standard deviation for the oxygen consumption rate of every sample across each stage.

##### NADH/NAD^+^ ratio quantification

All NADH autofluorescence signals were normalised to the OD_600_ measurement in the same well. The average of the technical triplicates that were included for each sample was used as the only value for that biological replicate. Afterwards, NAD pools were calculated by subtracting the reading acquired when all NAD molecules were assumed to be oxidised (NAD^+^) from the one at which all NAD was predominantly in its reduced state (NADH). Having estimated the NAD pool for each sample, we calculated the redox ratio of each well by discounting the background NADH fluorescence from the basal NADH measurements and dividing the resulting value over that of the NAD pool. This strategy was applied to obtain the redox ratios of three independent biological repeats per strain or cell line. Statistical analysis (unpaired t-tests) and visualisation were performed in GraphPad Prism 8.1.

##### Whole transcriptome analyses

Raw sequencing outputs were returned as FASTQ files. Hence, after conversion into FASTA format, files were processed using the paired-end (PE) version of the pyCRAC pipeline^10^. This package demultiplexes barcoded reads and removes adapter sequences by applying the previously developed Flexbar tool^55^. The resulting reads undergo two quality control steps. Firstly, sequences are filtered based on their length and nucleotide quality values (i.e., PHRED scores). Secondly, since PCR artefacts will display identical random prefixes, they can easily be pinpointed and excluded from downstream analysis (collapsed). The remaining unique reads were subsequently aligned to the corresponding regions of the Saccharomyces_cerevisiae.R64-1-1.75 reference genome with the Novoalign tool (version 2.07). The pyReadCounters function of the pyCRAC package (version 1.5.2)^10^ was used to compute the number of reads that map to genomic features. The resulting tables for all the samples were merged into a single TXT file that was analysed by the DESeq2 R package^24^. The pipeline corrects for size variability by computing a scaling factor for each sample^24^. To obtain such a normalisation factor, the software calculates the ratio of the raw count of a given gene in one of the samples to the geometric mean of the counts for that same gene across the compared samples^24^. The median of all the ratios calculated within a sample is defined as the scaling factor for that replicate^24^. Having adjusted the raw reads, the package then performs a logarithmic transformation of the data and calculates a fold change and an FDR for each entry^24^. Statistical significance was assessed for entries which exceeded or were lower than log2 fold change of 1 and −1 respectively^24^ and had an FDR value of 0.05 or less.

##### Proteomics analyses

Raw data were processed using MaxQuant version 1.6.5.0 with the Andromeda search engine^59,79^ for peptide identification. Inputs included a CSV file detailing the experimental design, a FASTA database (UP000002311 559292 YEAST Saccharomyces cerevisiae), and RAW files. Trypsin was used as the protease, with a minimum peptide length of 7 amino acids, allowing up to 2 missed cleavages. Initial peptide tolerance was set at 20 ppm, reduced to 4.5 ppm for main searches, with a peptide spectrum match (PSM) threshold of 1% FDR for statistical significance. Cysteine carbamidomethylation was a fixed modification, while methionine oxidation and protein N-terminal acetylation were variable modifications.

Post-analysis, MaxQuant outputted a proteinGroup TXT file, which was adjusted to include only intensity columns for comparison sets. This matrix was analysed using Perseus^62^ for statistical evaluation, filtering out PTMs, reverse proteins, and contaminants. Intensities were log-transformed, and proteins not detected in at least two replicates per strain were excluded. Missing values were imputed using a downshifted normal distribution. Statistical analysis of the log2-transformed data was conducted using EdgeR^54^, which performed differential expression analysis. Significant changes were defined by fold changes greater than 1 or less than −1, indicating a two-fold difference in expression relative to the parental strain.

##### Gene ontology analyses

Biological processes enrichment analysis was performed using PANTHER (Protein ANalysis THrough Evolutionary Relationships)^80^. For transcriptomic assessment, RNAs displaying a two-fold differential expression were used as input for the analysis. For proteomic evaluation, all significantly differential factors were fed to the PANTHER software. In the latter case, fold thresholds were omitted to provide a sufficiently large sample size from which the PANTHER suite could draw statistical conclusions on enriched GO terms. Bar plots were generated in GraphPad Prism 8.1.

